# Probing visual literacy skills reveals unexpected student conceptions of chromosomes

**DOI:** 10.1101/2024.09.20.614153

**Authors:** Crystal Uminski, Dina L. Newman, L. Kate Wright

## Abstract

Molecular biology can be challenging for undergraduate students because it requires visual literacy skills to interpret abstract representations of submicroscopic concepts, structures and processes. The Conceptual-Reasoning-Mode framework suggests that visual literacy relies on applying conceptual knowledge to appropriately reason with the different ways of representing concepts in molecular biology. We used this framework to specifically explore visual literacy related to chromosomes. We conducted 35 semi-structured interviews with students who had taken at least a year of college-level biology courses, and we asked them to sketch chromosomes, interpret an abstract representation of chromosomes, and use the abstract representation to answer a multiple-choice question about meiosis. While many participants used the correct vocabulary to describe chromosome structure and function, probing their visual literacy skills revealed gaps in their understanding. Notably, 97% of participants (34 of 35) held conceptual errors related to chromosome structure and function, which were often only revealed in their sketches or explanations of their sketches. Our findings highlight the importance of scaffolding visual literacy skills into instruction by teaching with a variety of visual models and engaging students in using and interpreting the conventions of abstract representations of chromosomes.

## INTRODUCTION

Molecular biology is often a difficult subject for students because the concepts in the curricula cannot be directly visualized (Johnstone, 1991; Newman et al., 2023). To make the invisible visible, biologists often create abstract models to represent molecular biology concepts. Abstract models are composed of shapes and symbols that highlight the salient structures and features that convey contextually important information (Fan et al., 2023). As such, abstract models are useful teaching tools in molecular biology because they allow instructors and students to focus on the most relevant information rather than extraneous or superfluous details. Biology instructors can use abstract models in their teaching to create meaningful opportunities for model-based reasoning that encourage students to deeply think about the purpose and efficacy of particular representations (Quillin & Thomas, 2015).

Abstract models are an important and necessary part of molecular biology education, but they can be a source of challenge or confusion for students, especially when considering how biologists often use different abstract shapes and symbols to emphasize varying aspects of the same concept (Kindfield, 1994b; Offerdahl et al., 2017; Schönborn & Anderson, 2006; Wright et al., 2022). For example, a biologist may choose to represent a gene as a sequence of letters to discuss a mutation, as boxes to point out the positions of exons, or as a double helix to explain the shape of the molecular structure. Given that there are a range of ways that biologists use abstraction in molecular biology, students need to develop skills to understand how, why, and when to use abstract models. The importance of these skills is underscored by their inclusion in the “Modeling” and “Communication and Collaboration” core competencies for undergraduate biology outlined in the *Vision and Change* report (American Association for the Advancement of Science, 2011; Clemmons et al., 2020a, 2020b).

The skills that biologists use to create and interpret abstract representations are also aligned with the broader set of skills associated with visual literacy. Visual literacy describes a viewer’s awareness of how conventions in visual representations convey meaning (Messaris & Moriarty, 2004), and definitions of visual literacy are typically refined to reflect the conventions and perspectives in a specific disciplinary context (Avgerinou & Pettersson, 2011; Trumbo, 1999). In molecular biology, visual literacy is often defined as the ability to “read” and “write with” the conventions and symbols that encode meaning in biochemical representations (Offerdahl et al., 2017; Schönborn & Anderson, 2006; Towns et al., 2012).

We can characterize the processes involved in visual literacy using the Conceptual-Reasoning-Mode (C-R-M) framework (Schönborn & Anderson, 2009, 2010). This framework proposes that accurately interpreting the many types or modes (M) of biochemical representations requires an understanding of the content knowledge relevant to the concepts being represented (C) and appropriately applying cognitive processes or reasoning (R) to perceive and decode visual conventions in the representation. These three factors (C, R, and M) are interactive, and the integration of all three factors is necessary for successful interpretation of a representation (Schönborn & Anderson, 2009). We provide an example of how the C-R-M framework can be applied to interpret an abstract representation of chromosomes in Figure 1.

**Figure 1.**
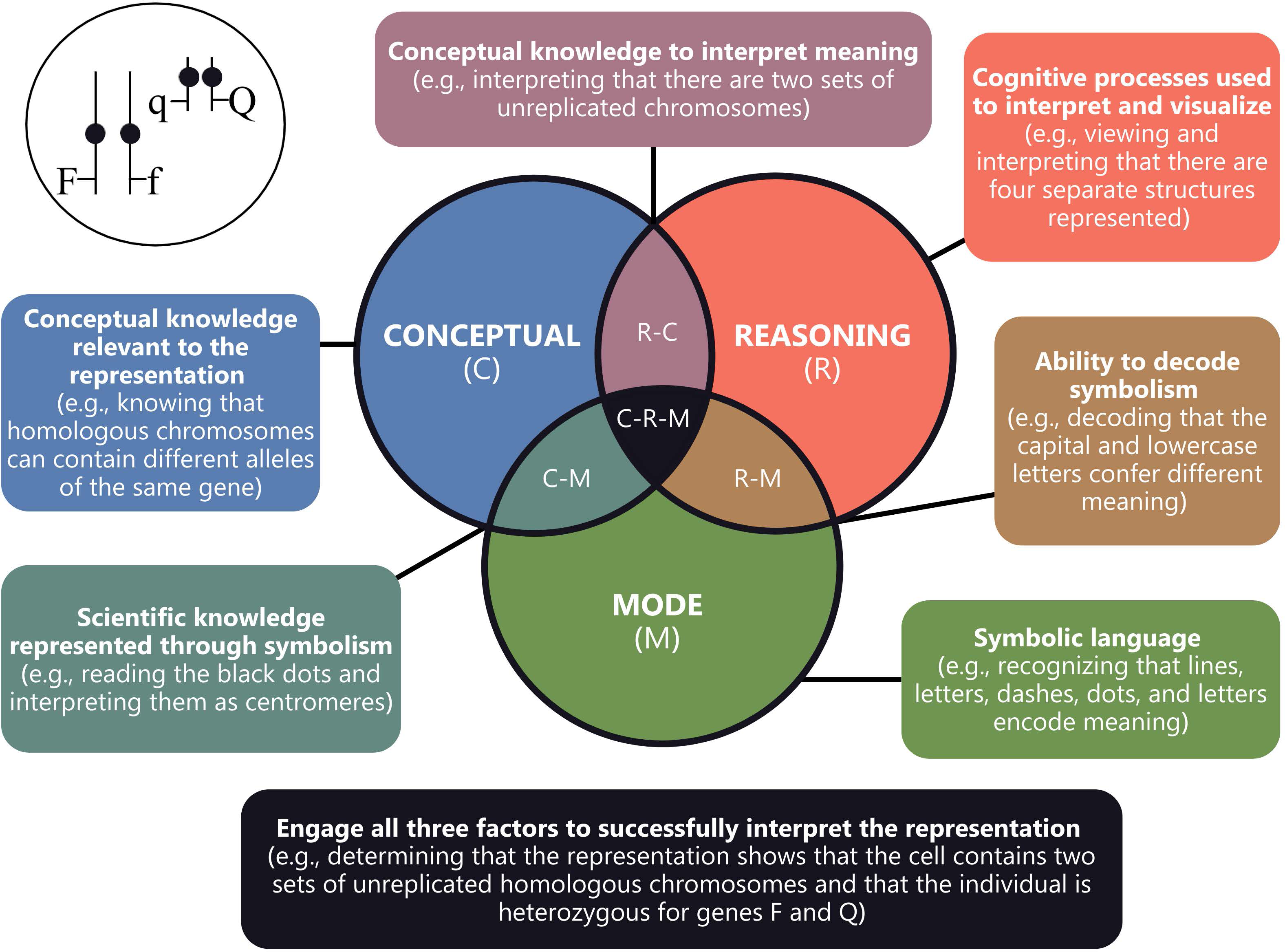
Application of the Conceptual-Reasoning-Mode framework to explain how a student might interpret an abstract representation of chromosomes. The representation of chromosomes in the top left is reproduced from item #2 in the Genetics Concept Assessment (Smith et al., 2008).

Expert-level visual literacy skills encompassed within the Reasoning factor of the C-R-M framework include the abilities to evaluate the power and limitations of representations, use representations to solve problems, and translate across multiple representations of a concept (Schönborn & Anderson, 2010). While these skills are necessary for reasoning and communication in molecular biology, they take time, practice, and scaffolded support to develop (Offerdahl et al., 2017). Experts develop visual literacy skills through their experiences interpreting, creating, and communicating with a breadth of visual representations (Arneson & Offerdahl, 2018; Kozma & Russell, 1997), so helping students develop expert-like visual literacy requires intentional allocation of class time for students to practice evaluating abstract representations, complete formative assessments using models to solve a problems, and draw a variety of representations of the same concept.

Instructors who create opportunities for students to develop visual literacy skills ought to be attentive to places where students may benefit from additional support. Previous research found that students may encounter challenges in maneuvering between different abstract representations of the same concept (Cho et al., 1985; Newman et al., 2023). Students may focus on the particular shapes, colors, and symbols that are used in a representation (i.e., the “surface features”) without necessarily decoding the meaning of the features in the representation (Newman et al., 2023). In some cases, such as in situations where students are using familiar scenarios and representations, finding patterns in the surface features may be a productive strategy, but solely relying on surface features may steer students away from understanding why a particular representation of a concept was chosen (Arneson & Offerdahl, 2018).

Instructors should also consider that students may need more structured support when they are using visual representations to reason across levels of organization or biological scale (Schönborn & Anderson, 2009). DNA is often a uniquely challenging topic for students to learn because the structure is both incredibly small and incredibly big in terms of the physical size and the number of base pairs, respectively. As such, biologists may choose to represent DNA at the scale of a nucleotide, a gene, or a chromosome, depending on what information they intend to communicate. Because there are multiple representations of nucleotides, genes, and chromosomes, the challenge of understanding visual conventions of scale can frequently be compounded with the challenges of abstraction.

We can better understand how students’ visual literacy in molecular biology is related to their conceptions of scale and abstraction in visual representations by applying a conceptual framework called the DNA Landscape (Wright et al., 2022). The DNA Landscape categorizes the different representations of DNA in a 3-by-3 matrix to show variation in representations across scale (nucleotide, gene, and chromosome) and abstraction (very abstract, elements of shape and abstraction, and literal shape). Students who are fluent in molecular biology visual literacy should be able to understand the relationships between images that are presented at different levels of abstraction and between representations at different scales, but to develop this fluency students must have previously productively engaged and appropriately reasoned with multiple representations at each scale (Schönborn & Anderson, 2006).

We focus here on how students conceptualize abstraction in representations of chromosomes. Conceptual knowledge of chromosomes is necessary for students to understand the *Vision and Change* core concept of Information Flow, Exchange, and Storage (American Association for the Advancement of Science, 2011; Brownell et al., 2014), but historically students often struggle with course content related to chromosomes (Etobro & Banjoko, 2017; Kindfield, 1991; Newman et al., 2012; Saka et al., 2006). Chromosome structure can be a particularly difficult topic for students (Kindfield, 1991, 1994a, 1994b; Newman et al., 2012; Wright et al., 2021), and misunderstandings related to the function and behavior of chromosomes in mitosis and meiosis are so common among students that these concepts are targeted on concept assessments such as the Introductory Molecular and Cell Biology Concept Assessment (Shi et al., 2010), the Genetics Concept Assessment (Smith et al., 2008), and the Meiosis Concept Inventory (Kalas et al., 2013). Although these misunderstandings are commonly identified in the literature, little work has been done to examine the relationship between how students’ conceptualization of chromosomes may be associated with their visual literacy skills in molecular biology (Newman et al., 2023). Thus, we set out to answer the following research question: How do students conceptualize chromosomes and apply visual literacy skills to interpret abstract images of chromosomes?

## METHODS

### Participants

We conducted semi-structured interviews with 35 students from two different institutions—a primarily undergraduate institution and an R2 research institution (Table 1). All research participants had completed at least a year of biology coursework. We used several recruitment methods to achieve a broad sample of participants, including emailing biology majors at each institution, posting flyers in science buildings, and soliciting volunteers in person during sessions of upper-division biology courses. We also used snowball sampling by asking participants to refer their peers to register for the study after they had concluded the interview protocol. Participants consented to this research and were compensated with a $20 Amazon gift card for their participation. Pseudonyms were selected by the participant or assigned by the researcher. This research was approved and classified as exempt from human-subjects review by Rochester Institute of Technology (protocol 01090823).

**Table 1.**
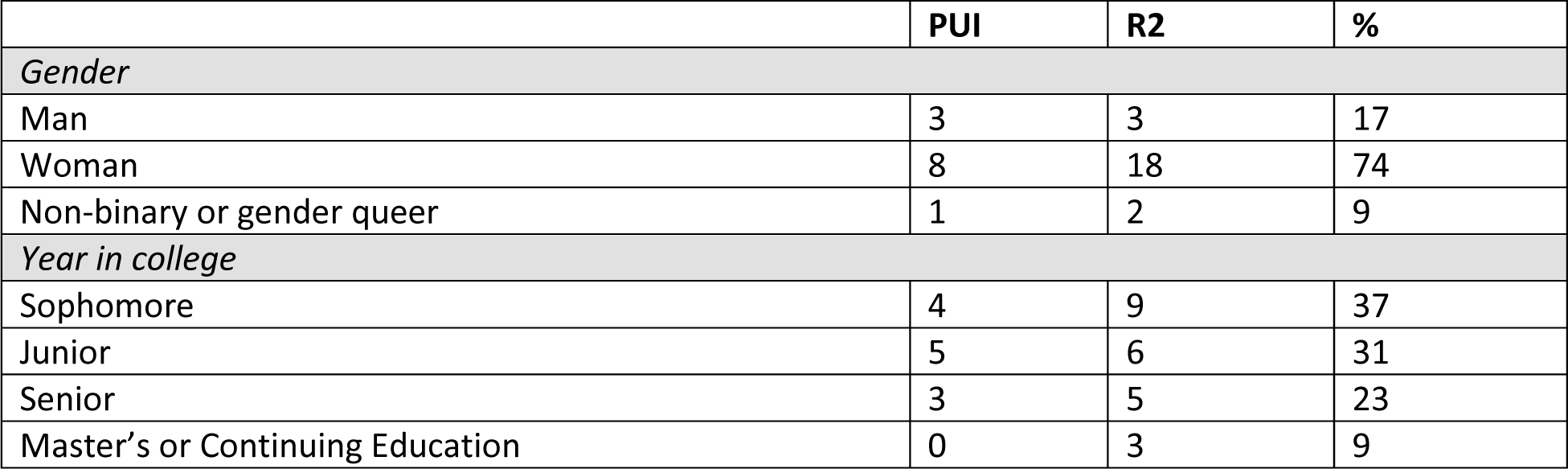
Research participant demographics.

| Table 1. Research participant demographics |  |  |  |
| --- | --- | --- | --- |
|  | PUI | R2 | % |
| *Gender* |  |  |  |
| Man | 3 | 3 | 17 |
| Woman | 8 | 18 | 74 |
| Non-binary or gender queer | 1 | 2 | 9 |
| *Year in college* |  |  |  |
| Sophomore | 4 | 9 | 37 |
| Junior | 5 | 6 | 31 |
| Senior | 3 | 5 | 23 |
| Master's or Continuing Education | 0 | 3 | 9 |

### Interview Protocol

We developed the prompts in our interview protocol to align with two frameworks to target specific visual literacy skills (Table 2). First, we designed interview prompts to target the factors from the Conceptual-Reasoning-Mode (C-R-M) framework that are associated with the ability to interpret and visualize biological representations (Schönborn & Anderson, 2009, 2010; Figure 1). We wrote our interview prompts in ways that could elicit evidence of participants’ conceptual understanding of chromosomes (Conceptual) as well as the cognitive skills participants engaged (Reasoning) to make sense of visual representations (Mode) of chromosomes. The C-R-M framework indicates that there is a minimum amount of prior conceptual knowledge that is necessary for the interpretation of visual representations (Schönborn & Anderson, 2010), so we scaffolded our interview questions to target conceptual understanding before asking participants to reason about the visual representation we presented in the interview. Second, we refined our use of the Reasoning component of the C-R-M model by classifying cognitive skills according to Bloom’s Taxonomy (Anderson et al., 2001; Bloom et al., 1956). We used the Visualization Blooming Tool (Arneson & Offerdahl, 2018) to design interview prompts that target visual literacy skills across a range of Bloom’s Taxonomy cognitive skills. As with any prompts or questions designed to target specific cognitive skills, the mental processes that a participant engages may not always align with the targeted cognitive skill level (Monrad et al., 2021).

**Table 2.** Interview protocol and alignment of interview prompts to visual literacy frameworks.

|  |  |  |
| --- | --- | --- |
| Interview Prompt | C-R-M<sup>a</sup> | Bloom's Level(s)<sup>b</sup> |
| "What is a chromosome?" | C | Knowledge |
| "Can you sketch a chromosome?" | C-M | Knowledge |
| "Why did you draw the chromosome this way?" | C-R-M | Knowledge/<br>Comprehension |
| Participant is presented with the image of chromosomes from Genetics Concept Assessment item #2.<br>"Please explain what you think about this representation of chromosomes." | C-R-M | Comprehension |
| "What about this image makes this an effective representation of chromosomes?" | C-R-M | Evaluation |
| "Is there anything surprising or confusing about this representation of chromosomes?" | C-R-M | Evaluation |
| "How does your sketch of a chromosome compare to this representation? What is similar and what is different between your sketch and this image?" | C-R-M | Analysis |
| Participant is presented with the text from the GCA item #2. "Please talk through how you answered this question." | C-R-M | Comprehension/<br>Application |
| "Can you sketch a representation of the answer you selected?" | C-R-M | Application |
| *Notes*<br><sup>a</sup>Abbreviations: C = Conceptual; R = Reasoning; M = Mode (Schönborn & Anderson, 2009)<br><sup>b</sup>Bloom's levels classified based on the Visualization Blooming Tool (Arneson & Offerdahl, 2018) |  |  |

We incorporated an item from the validated and widely-used Genetics Concept Assessment (GCA; Smith et al., 2008) into our interview protocol. Items that compose concept assessments like the GCA are typically evaluated for statistical measures that indicate item difficulty (i.e., the number of correct responses divided by the total number of responses) and item discrimination (i.e., the ability of the item to distinguish between high- and low-scoring students). We used item #2 from the GCA, which had posttest item difficulty of approximately 0.7 and item discrimination index of approximately 0.4 (Smith et al., 2008). These values indicate that this item is sufficiently difficult and can effectively discriminate between students with high and low test scores.

Our interview protocol (Table 2) used a scaffolded structure to prompt research participants to engage in higher-order cognitive skills. We first asked participants to provide a verbal definition of chromosomes and then to draw a chromosome and explain their drawing. Some participants drew additional sketches to supplement their explanations. Next we prompted participants to interpret the illustration that accompanied item #2 from the GCA. The GCA item includes an abstract representation of two pairs of unreplicated homologous chromosomes within a cell (see reproduction of this representation in Figure 1). In this representation, the chromosomes are depicted as single vertical lines with large black dots for centromeres, and alleles on the chromosomes are represented using letters (F,f, Q, q). We asked participants to evaluate the effectiveness of the representation of chromosomes and to describe any confusing elements within the representation. Then we asked them to compare and contrast their drawing to the GCA representation. We next provided participants with the full item #2 from the GCA. The item includes a written description of the image that explains that it contains two pairs of chromosomes on which are two different genes, and it asks them to select the possible sperm genotypes if the cell divides normally. After participants selected their answer from a multiple-choice list, we asked them to draw a picture that represents the option they selected.

We conducted and recorded interviews using the video conferencing software Zoom. Of the 35 interviews, 11 participants (31%) created drawings using the Zoom whiteboard and 24 participants (69%) drew on their own paper and sent us photos or scans of their sketches via email. The quality of drawings and sketches on the Zoom whiteboards may have varied depending on the familiarity and comfort with the virtual medium. Audio recordings were transcribed by Otter.ai and then cleaned manually.

This interview protocol was part of a larger interview probing visual literacy across a range of molecular biology concepts. Participants completed the chromosome protocol in approximately ten minutes, while the entire interview was approximately 45 minutes in length.

After 35 interviews, the emergence of new insights or concepts from interviews had slowed, so we judged that we had reached a theoretical saturation of conceptions and reasoning about chromosome structure.

### Codebook development

We used initial coding (Saldaña, 2013) to develop our codebook from the interview data. We shared our initial codebook with a group of biology education researchers consisting of two biology professors, two postdoctoral researchers, and four undergraduate students. The group of researchers used the codebook to code one interview transcript that was selected based on a random-number generator. We compared interpretations of the codebook across the members of the research group, discussed areas of disagreement, and subsequently refined our codebook to its present version.

Our codebook consists of sets of codes to categorize participants’ verbal definitions of chromosomes, sketches of chromosomes, the accuracy of their description of their sketch, the accuracy of their interpretation of the concept assessment image, the accuracy of their answer to the concept assessment item, and the sketches that represent their response to the concept assessment item.

*Verbal definitions of chromosomes:* We coded verbal descriptions of chromosomes based on whether they described the shape (e.g. “X-shaped”), structure (e.g. “made of DNA”), and/or function (e.g. “involved in meiosis”) of chromosomes.

*Sketches of chromosomes*: We categorized sketches of chromosomes based on both shape and the style representing the curvature or complexity of the shape. Participants drew chromosomes as either an X-shape (consisting of two structures that cross or intersect at a specific point that may or may not have a visible centromere indicated), an I-shape (a single vertical structure that may or may not have a centromere indicated), or as a side-by-side shape (two structures that do not intersect). For style, participants drew chromosomes consisting of loops (a rounded, continuous curve that creates an enclosed shape), lines (a single continuous pen stroke without additional details), or coils (tightly-would spirals or twists). We also coded sketches for their level of abstraction based on their location on the DNA Landscape matrix (Wright et al., 2022 *Accuracy of description of sketch*: We classified participants’ descriptions of their sketches as either correct (participants used the correct vocabulary to describe the elements in their sketch, like correctly stating that their X-shaped chromosome was a replicated chromosome) or incorrect (participants used incorrect vocabulary or concepts to describe their sketch, like identifying an X-shaped chromosome sketch as a pair of homologous chromosomes).

*Concept assessment image interpretation*: We classified participants’ interpretation of the image from the GCA item #2 as either correct (stated the image consisted of four chromosomes or two pairs of homologous chromosomes and that the letters in the image represented genes or alleles) or incorrect (stated the number of chromosomes was something other than four, mistook the homologous chromosomes for sister chromatids, or said the homologous chromosomes should be connected at the centromere).

*Answer to concept assessment*: We coded three types of answers to the GCA item #2: the correct answer with the correct reasoning, the correct answer with incorrect reasoning, or an incorrect answer.

### Interrater reliability

To establish interrater reliability between coders, two members of the research team (CU and LKW) used the codebook to independently code eight randomly selected interview transcripts (23% of the sample). There was 93% agreement between the two raters across 160 coding decisions. The Cohen’s kappa coefficient was 0.83, which indicates strong agreement between raters (McHugh, 2012). The two raters discussed any disagreements until consensus.

### Positionality

Author CU conducted the interviews and acknowledges the limitations in this research because of the power differential inherent in their position as an education researcher who is older and has more formal education training relative to the interviewees. This power differential may inadvertently reinforce traditional power structures and hierarchies, thereby shaping the ways in which participants perceived and responded to the interview.

All members of the research team are trained in biology education research, have studied visual representations in biology, and have taught courses where we observed students making errors about chromosome structure and/or function. As such, we had preexisting ideas about how research participants might respond which consciously and unconsciously shaped our analysis and interpretation of participant responses.

## RESULTS

We conducted 35 semi-structured interviews to understand how students conceptualize chromosomes. The participants in our study brought with them a wealth of experiences in biology that allowed them to thoughtfully engage in interpreting, evaluating, and creating abstract representations of chromosomes. While it was not directly part of our structured interview protocol to ask students where they previously encountered representations of chromosomes, many participants volunteered information about their experiences learning about chromosomes in high school (n = 2), in their undergraduate courses in introductory biology (n = 3), molecular biology (n = 1), genetics (n = 5), as well as in biology classes or lectures in a broader sense (n = 9). In addition to their courses as sources of information, participants discussed seeing chromosomes represented in textbooks (n = 5), microscopy (n = 5), and karyotypes (n = 2). Although many participants openly discussed their experiences learning about chromosomes, our results as a whole indicate that most participants likely left these experiences with an incomplete understanding of the representations they had encountered.

When we look across the entirety of participant’s interview, we found that 97% of participants (34 of 35) demonstrated at least one error in their conceptual knowledge of chromosomes. Using the Conceptual-Reasoning-Mode (C-R-M) framework as a lens for our research, we frequently found that most participants knew the correct vocabulary to describe chromosomes (C) and were familiar with the shapes and symbols used in common abstract models of chromosomes (M). However, we observed mismatches between participants’ verbal statements and their drawings that indicated an incomplete conceptual understanding of why and how expert biologists typically use abstract shapes and symbols to convey meaning (C-M). We further probed participants’ understanding of chromosomes using an item from the Genetics Concept Assessment and found participants often had difficulties reasoning with the unfamiliar chromosome representation in the item (R-M). Despite misinterpreting the GCA image, the majority of participants correctly answered the associated concept assessment item; however, when asked to explain their thinking, most participants provided an incorrect line of reasoning to justify their answer.

### How do undergraduate students define chromosomes?

To investigate what conceptual knowledge of chromosomes participants were bringing into the interview (the C in C-R-M), we first asked them “What is a chromosome?”. The majority of participants (86%, n = 30) provided definitions related to chromosome structure (Figure 2). Most participants correctly noted structural elements of chromosomes, such as the fact that chromosomes consist of DNA and contain histones. Participants less frequently mentioned chromosome functions (e.g., the role of chromosomes in gene expression or meiosis) or chromosome shape (e.g., an “X” shape), but when they did mention function or shape, it was typically paired with content related to chromosome structure.

**Figure 2.**
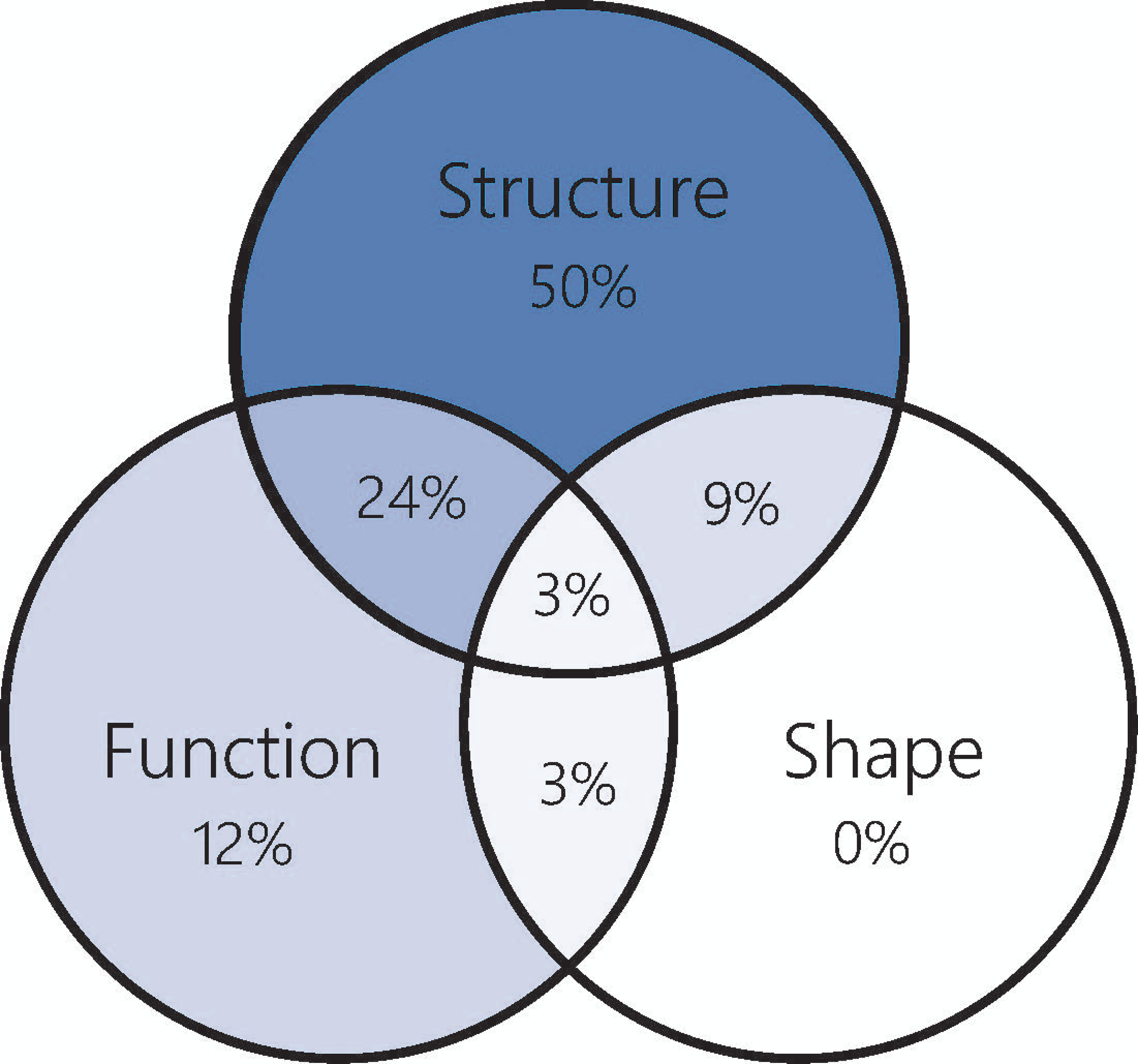
Percent of participants defining chromosomes based on structure, function, or shape. Overlapping regions indicate definitions that met two criteria, with the central overlap indicating a definition met criteria for chromosome structure, function, and shape. This Venn diagram uses blue color to indicate that definitions are located in the Conceptual factor of the C-R-M framework as shown in Figure 1.

While participants frequently provided factually correct definitions of chromosomes, we noted several unrefined ideas, often relating to the size and scale of chromosomes. For example, Jimmy (sophomore biotechnology major) stated, “A chromosome is I guess like a collection of DNA and genes that are—they’re like the encapsulation of all your DNA.” Jimmy inaccurately implies that a single chromosome contains all the DNA within what is presumed to be a eukaryotic cell. In other cases, participants confused the scale of chromosomes and genes. For example, Evelyn (junior biology major) said, “A chromosome is part of the DNA that can lead to a gene,” and Mia (sophomore biology major) said of chromosomes, “basically I think it’s like part of the genes.”

Additionally, we uncovered inaccurate ideas related to chromosome function during cell division. For example, Norma (senior biotechnology major) said, “A chromosome is basically like an X. It can be single, it can also be like the X shape, and it contains genetic information and just I guess, a shaped form. And the DNA can be double stranded—is double stranded—and then if it goes through like the other processes, it could become single stranded and whatnot.” While Norma understood that genetic information is divided in “other processes,” such as mitosis and meiosis, division does not make DNA into a single-stranded molecule.

### How do undergraduate students draw chromosomes?

We asked participants to draw a chromosome to explore how they used shapes and symbols to represent features of chromosomes that convey biological meaning (the M in C-R-M). When asked to sketch a chromosome, 100% of participants drew images that aligned to the “very abstract” representation of chromosomes on the DNA Landscape and drew symbolic loops, lines, or coils to represent X-shaped, I-shaped, or side-by-side chromosomes (Figure 3).

**Figure 3.**
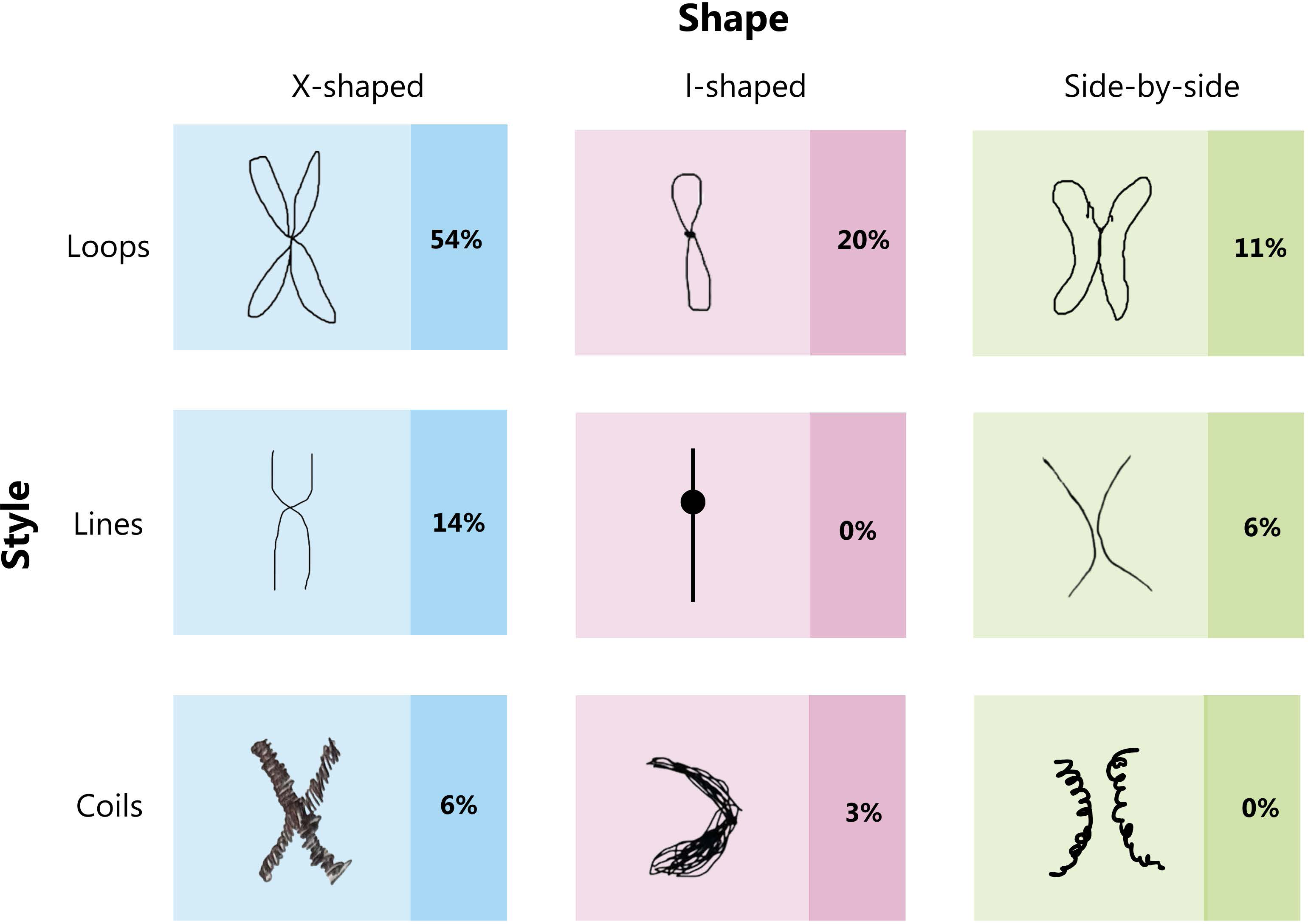
Categorization of participants’ initial sketches of chromosomes. Sketches of chromosomes are categorized by shape (X-shaped, I-shaped, side-by-side) and symbolic style (loops, lines, coils). Examples of each representation type are reproduced from participant sketches when available. From left to right, example sketches were from interviews with the participants Evelyn, Catherine, Mia, Donna, Christin, Jack and Ristan. There were no sketches for I-shaped lines or side-by-side coils. Some participants produced several different sketches in response to the interview prompt, so the sum of percentages exceeds 100%.

We found that participants most commonly drew X-shaped chromosomes (74%) and most frequently represented these X-shaped chromosomes using loops. Participants less commonly drew I-shaped (23%) and side-by-side (17%) chromosome structures. We did not observe any participants drawing I-shaped chromosomes with lines in ways that resembled the representational style used in the Genetics Concept Assessment image.

### Why do students draw chromosomes that way?

To understand how participants were representing their conceptual knowledge of chromosomes in their sketches, we asked them to explain the reasons why they drew chromosomes in the way that they did. We group these explanations based on the shape and style of the chromosome sketch.

*X-shaped loops*: We found that X-shaped loops were the most frequently drawn representation. In some cases, participants focused the explanation of their sketches on their prior experiences with similar representations. “Well, that’s just how I’ve always seen it in any of my classes—it’s always two sister chromatids held together by a centromere,” Maddy (junior biology major). For participants like Maddy, their description aligns more closely with how an expert might describe and use this representational form.

For other participants, their description of the meaning of the X-shaped loops was more ambiguous or did not entirely align with how experts conventionally use this mode of representing chromosomes (Figure 4). For example, Nadine (senior environmental science major) describes her X-shaped loops as consisting of “two halves that kind of fit together,” but it is unclear how or by what process she thought that this “fit” occurs. In another example, Annie (sophomore biochemistry and molecular biology major) described that her sketch contained “four parts” across two sections of genetic code (Figure 4). When the interviewer asked for clarification about this statement and whether the “two sections” (i.e., the two p-arms in a replicated chromosome) that Annie referenced were identical, she responded “Um, no. So like, for example, if you have like, if you are homozygous it’d be like your big E and then like the little e like that.” Here, Annie is not only uncertain about the correct vocabulary (i.e., homozygous compared to heterozygous), but conflated sister chromatids (which must contain the same allele) in her drawing with homologous chromosomes (which could have two different alleles).

**Figure 4.**
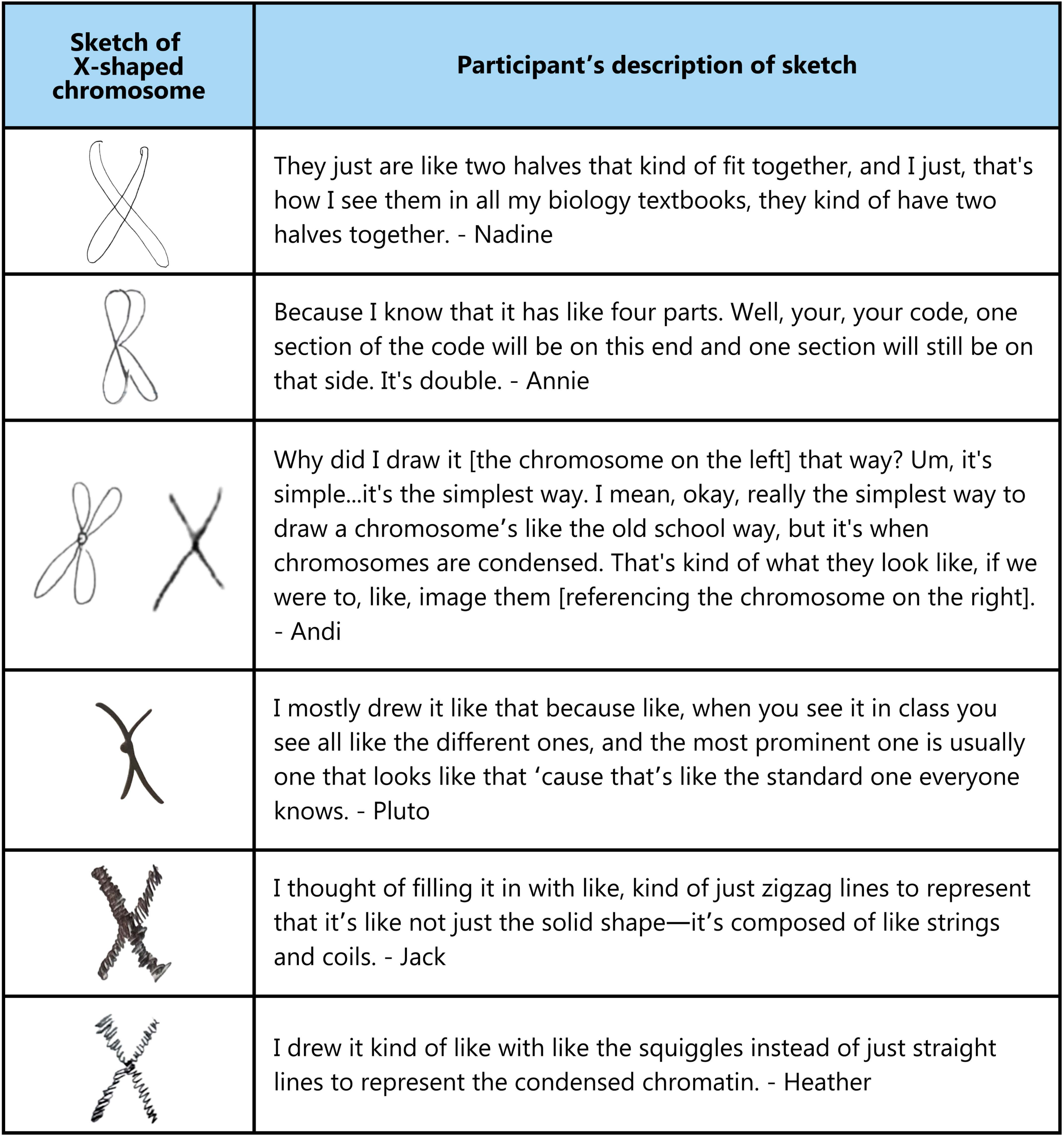
Initial sketches of X-shaped chromosomes and the associated verbal description of the sketch. Nadine, Annie, and Andi drew chromosomes with X-shaped loops. Andi and Pluto drew chromosomes with X-shaped lines. Jack and Heather drew chromosomes with X-shaped coils.

We found that some participants used the X-shaped loops to productively reason about chromosome structure. Evelyn (junior biology major), whose sketch is the exemplar of the dimensional X-shape in Figure 3, described it as a “circular thing [so] that you could probably see that there is more like their stuff inside it.” Here, the loops are meaningful symbols that reflect her understanding that chromosomes are structures that are composed of “stuff,” which we interpret as DNA and proteins.

*X-shaped lines*: Participants less commonly represented chromosomes using two crossed lines in an X-shape (Figure 4). Even though participants like Pluto (sophomore general science major) asserted that “everyone knows” the X-shaped line style of representing chromosomes, only 14% of participants chose to use this representation in their drawing.

*X-shaped coils*: We identified two instances of X-shaped coils which seemed to contain both elements of both the “very abstract” and “very literal” portions of the DNA Landscape (Figure 4). These sketches were both X-shaped but used coils as a way of representing the more realistic literal structure of chromosomes at the microscopic level. Jack (senior mechanical engineering major) rationalized that his coiled chromosome could be “filled,” which was similar to how participants used X-shaped loops to represent how chromosomes could have “stuff inside.”

*I-shaped loops*: Nine participants drew sketches of I-shaped loops. Two participants referred back to the interview prompt, “Can you sketch a chromosome?,” in their explanations. Josie (junior biochemistry and molecular biology major) asked the interviewer “Should I do like — I’ll just do one?” and proceeded to draw what we interpreted as a single unreplicated chromosome using I-shaped loops. Similarly, Lacie (junior biotechnology and molecular bioscience major) referred back to the interview question, “’cause I was like ‘just one chromosome.’” Both Josie and Lacie mentioned their sketches were not reflecting what they “normally” thought about for chromosomes. Josie remarked that “The way I normally picture them is double. Because we talked about them with like, the replication, but that’s—I’m pretty sure they just exist as single.” Lacie said that “I was thinking, you know, normally in human beings, chromosomes are drawn in pairs.” Neither of these participants used the terms “replicated” or “unreplicated” or discussed the total number of DNA molecules when deciding which type of chromosome structure to draw.

*I-shaped lines*: We did not observe any participants using I-shaped lines in their sketches of chromosomes. The lack of this representational style in the initial drawings tracks closely with the first-impressions that participants had to the I-shaped line chromosomes in the GCA image. For example, Donna (sophomore biology and environmental science major) viewed the GCA image and immediately stated “If I saw this, I would not recognize it as a chromosome.”

*I-shaped coils*: Ristan (senior biochemistry and molecular biology major) was the only participant to draw a chromosome using I-shaped coils (see Ristan’s sketch as the exemplar for I-shaped coils in Figure 3). He initially said “I think that’s to sort of show that it’s not like one strand.” Ristan provided a second sketch of a single line for comparison to better illustrate that his first sketch contained “many, many strands of DNA all compressed on top of each other,” suggesting that numerous individual molecules of DNA make up an unreplicated chromosome.

Participants sometimes drew coiled chromosomes during follow-up questioning which provided unique insights into their understanding of chromosome structure (Figure 5). These sketches are not included in the initial counts in Figure 3, but they provide contextual information about how students are illustrating meaning with I-shape coils. In Figure 5, we see that Norma’s initial sketches include X- and I-shaped loops, stating that a chromosome “becomes an X-shape throughout the process of meiosis.” When probed further, Norma produced the follow-up sketch and said “DNA is just coiled up, so it’s like this,” suggesting a robust understanding of chromosome structure. Conversely, Teresa (junior individualized study major) created follow-up sketches that indicate a *misunderstanding* of chromosome structure. Teresa modified her sketch by filling in the p-arm of the chromosome with the narration that “This is like the clusters of the DNA, big clusters.” She stopped at the centromere then continued in a separate pen stroke to fill in the q arm of the chromosome, saying “Yeah, and this is also another cluster of like DNA.” Her description shows that she thought of chromosomes as consisting of two separate units of DNA partitioned by the centromere.

**Figure 5.**
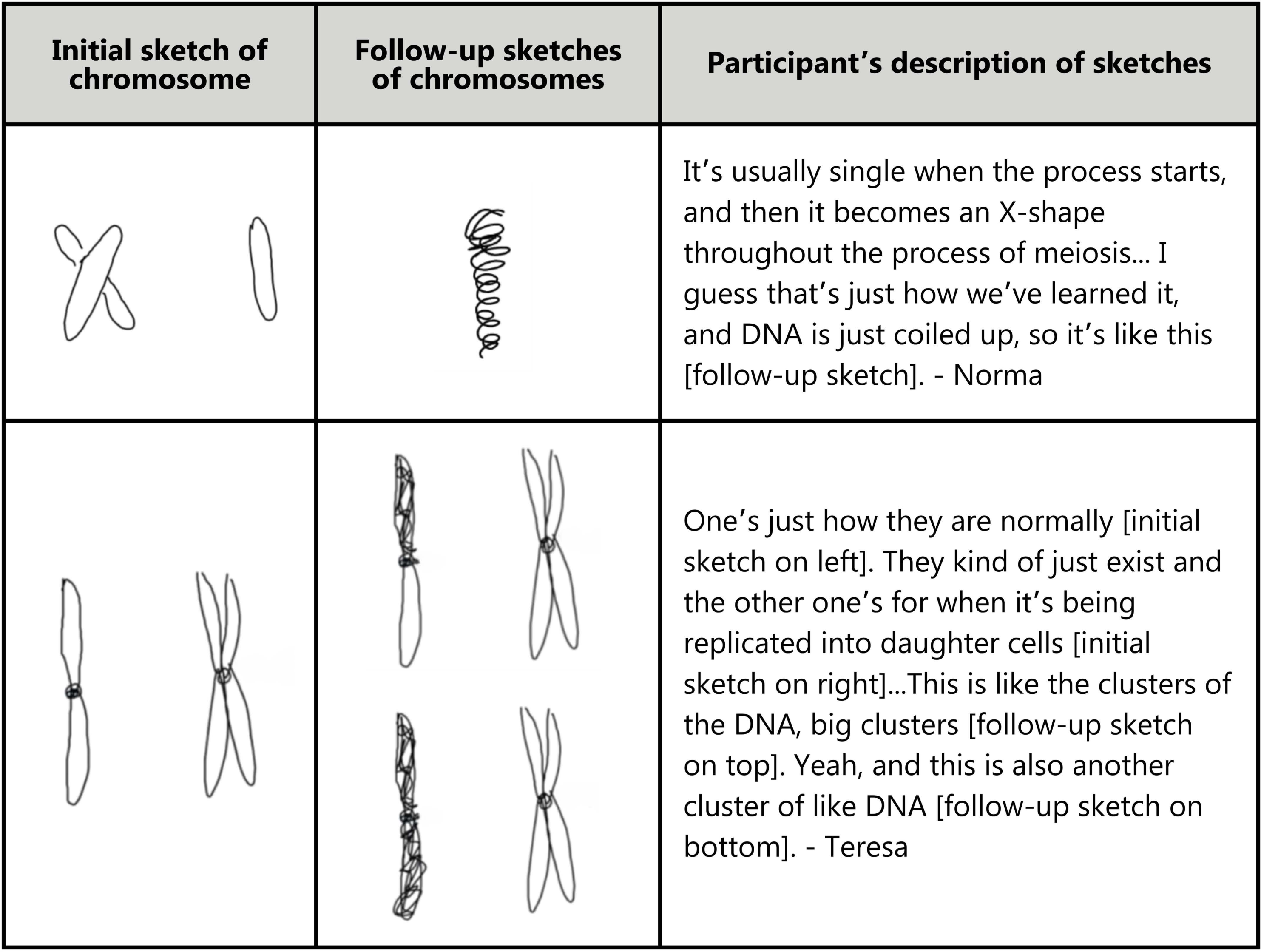
Participant sketches of coiled chromosomes generated during follow-up questioning. Participants’ initial sketches are in the first column. The second column shows the follow-up sketches. Note that in Teresa’s follow-up sketch, she modified her original drawing.

*Side-by-side loops*: There was variability in the participants’ rationale for their side-by-side chromosomes (Figure 6). Alex (senior biology major) drew the side-by-side chromosome with a separate box to represent the centromere, inappropriately indicating that the centromere is a structure separate from the structure of the chromatids. Mia (sophomore biology major) said that she drew side-by-side chromosomes to resemble X and Y chromosomes. Participants Socks (sophomore biomedical science major) and Leo (junior biology pre-med major) both used side-by-side representations, but offered very different explanations as to why. Socks said they drew side-by-side loops “because that’s usually how we see it on a karyotype—they’re usually paired together like that for mitosis,” which we interpreted as a description of unreplicated homologous chromosomes. While homologous chromosomes are typically shown paired in a karyotype, this participant is mistaken because homologous chromosomes do not pair for mitosis. This quote is also interesting because chromosomes in karyotypes are replicated (but sister chromatids are not visible as separate entities), so Socks is likely interpreting the chromosomes in karyotypes as single DNA molecules even though they are not. Leo, on the other hand, said “that would be a set of sister chromatids.” We highlight Socks and Leo as two examples of how the same sketch of a chromosome can be ambiguous if a verbal or written description is not provided.

**Figure 6.**
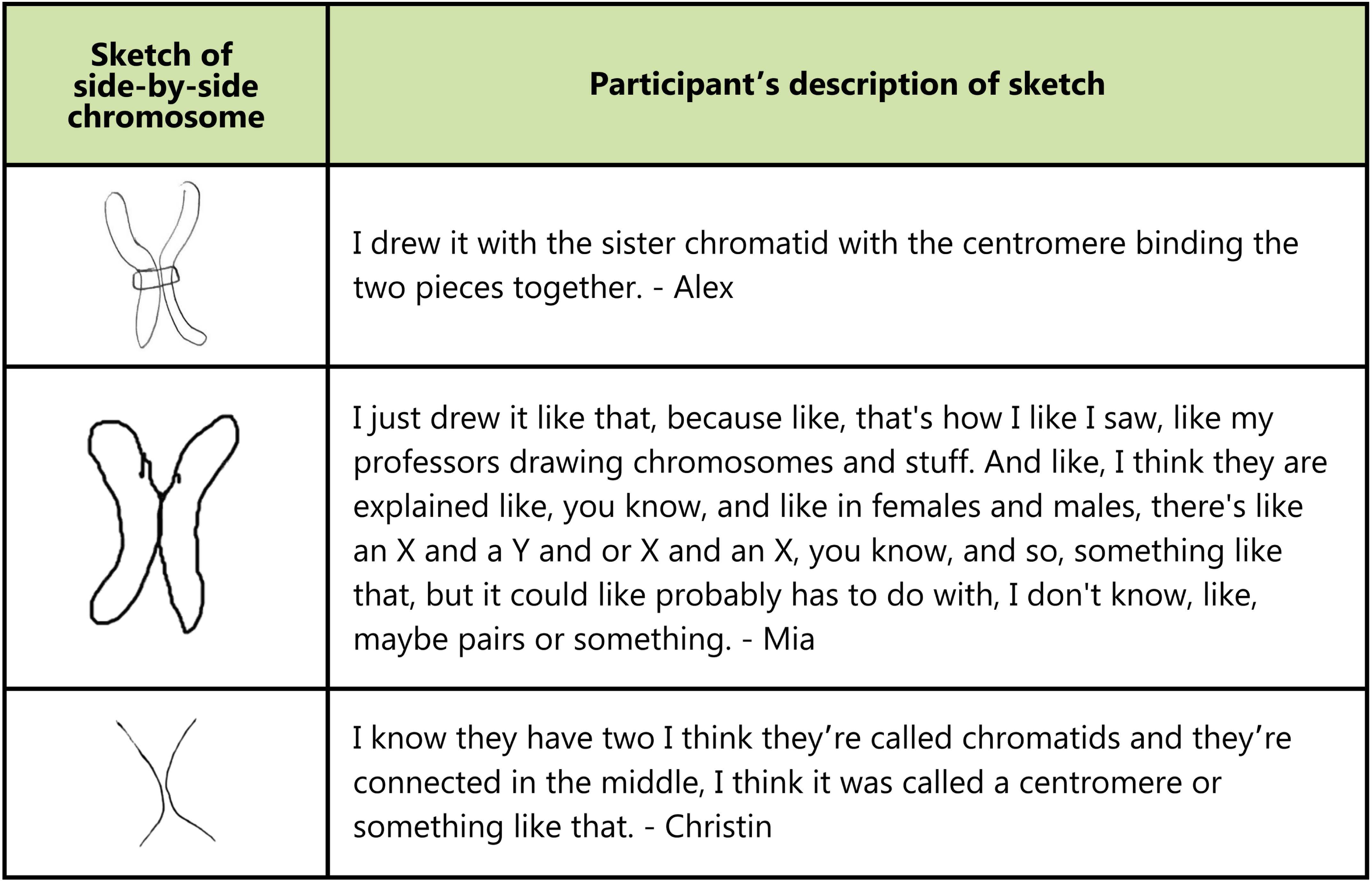
Initial sketches of side-by-side chromosomes and the associated verbal description of the sketch. Alex and Mia drew chromosomes with side-by-side loops, and Christin drew side-by-side lines.

*Side-by-side lines*: Christin (sophomore environmental science major) and Regina (post-baccalaureate continuing education) drew chromosomes with side-by-side lines, which they used to indicate different numbers of chromosomes. Christin referenced a centromere in her explanation of her drawing, even though her sketch did not include a centromere, suggesting that she intended the two lines to represent chromatids in a single replicated chromosome (Figure 6). Regina said “When I think of chromosomes, I think of mitosis and meiosis, and they’re always paired up with their buddies.” To clarify this explanation, Regina drew an arrow pointing to only one of the lines in the sketch and indicated that there were two chromosomes but “they’re like a pair.” While Christin used the lines to represent one chromosome, Regina used the same style of lines to represent two.

While we focus on what participants did draw in their sketches of chromosomes, it is also important to note what they did not draw. None of the participants in our sample drew chromosomes in a circular shape that might resemble bacterial chromosomes. We did not observe any participants drawing chromosomes with banding patterns like a karyotype or drawing a map with positions of genes.

We necessarily focus our results on the errors related to chromosomes, as these errors provide novel insights into student thinking, but we also want to emphasize many participants (63%, n = 22) used appropriate terms and vocabulary to describe the chromosome structure that they drew.

### How do undergraduate students interpret an abstract model of chromosomes?

To determine how participants use their conceptual knowledge to reason about an abstract model of chromosomes, we asked them to interpret an image from the Genetics Concept Assessment. To correctly interpret the image, participants needed to appropriately engage all three factors in C-R-M and apply their conceptual knowledge (Figure 1). While many students correctly decoded certain symbols, such as the capital and lowercase letters representing dominant and recessive alleles, students may have had incomplete content knowledge and/or an unfamiliarity with the abstract image that prevented them from accurately interpreting all of the key information encoded in the image. Overall, we found that 71% of participants (n = 25) incorrectly interpreted the image from the concept assessment (Table 3).

**Table 3.** Participant interpretation and responses to an item from the Genetics Concept Assessment.

|  |  |  |  |
| --- | --- | --- | --- |
|  | Incorrect interpretation of image | Correct interpretation of image | Total |
| Correct response to item with correct reasoning | 3 | 5 | 8 |
| Correct response to item with incorrect reasoning | 16 | 2 | 18 |
| Incorrect response to item | 6 | 3 | 9 |
| Total | 25 | 10 | 35 |

For some participants, accurate interpretation of the GCA image was most directly limited by an unfamiliarity with the mode of representation. G (sophomore biotechnology major) and Donna (sophomore biology and environmental science major) compared the unfamiliar representation in the GCA to the X-shapes they were more familiar with. For example, Donna said “I’m used to seeing chromosomes where they’re crossed over and in an X formation, so that’s kind of the only reason I wouldn’t recognize this because it doesn’t have the X formation.”

Donna and G also held conceptual misunderstandings about the difference between homologous chromosomes and sister chromatids. Upon seeing the GCA image, G immediately asked “The chromatids aren’t paired together?”. G subsequently described the image as having “all the information needed—it has two different chromatids.” These types of conceptual misunderstandings—thinking that sister chromatids can exist independently or that parental origins are mixed in a replicated chromosome—are misunderstandings that underpinned the majority of the misinterpretations of the GCA image, occurring in 19 of the 25 incorrect interpretations.

This confusion of homologous chromosomes and sister chromatids persisted even with the visual scaffolding provided by the heterozygous alleles labeled for each chromosome pair (F/f and Q/q). Maddy (junior biology major) incorrectly used the alleles to justify her interpretation of the image as chromatids, stating that the labeled alleles are “just like to specify that they’re the same chromosome but different sister chromatids.” Heather (sophomore biotechnology and molecular bioscience major) similarly struggled to differentiate chromosomes and chromatids, stating that “in terms of chromosomes that have already been replicated, there’s two. So that would be a total of four.” In other words, Heather arrived at the correct number of chromosomes in the image by applying the wrong reasoning.

In many cases, this misunderstanding of homologous chromosomes stemmed from the idea that the unreplicated homologous chromosomes in the image were “separated” sister chromatids. Participants tended to view the centromere as the place where the chromatids “should have been paired together” or connected. Numerous quotes illustrate this point:

“The two halves of the chromosome just pulled apart—Instead of being in that X, they’re just portrayed separately” – Eileen (sophomore environmental science major) “I usually imagine [chromosomes] like connected like I did [in my sketch of a replicated chromosome]. Because they’re like separate—might be something that I thought was a little confusing at first.” – Quentin (junior biology/pre-med major) “[The image is] like showing that there’s two separate chromatids, and because like on the initial drawing that I drew [this participant initially drew a replicated chromosome], a lot of people would just see it as like one X structure and not like two structures that are together.” – Jimmy (sophomore biotechnology major)

“In both sketches [participant sketch and the GCA image], it’s made out of two parts and you can kind of see that they’re supposed to be attached together with the dots on that sketch and on my sketch.” – Sylvia (sophomore biochemistry and molecular biology major)

“Typically, I think they’re [the chromatids] kind of fused together, so maybe that’s one thing that might be confusing to people [who] just started learning.” – Jay (Master’s student in bioinformatics)

Some participants misapplied conceptual knowledge in how they reasoned about the length of the chromosomes in the image. For example, Annie conflated the length of the chromosomes with dominance in alleles: “Could the longer strands mean it’s dominant?” Brittany (senior biology major), interpreted the different lengths as being related to a mutation. “My first thought was that there was like a weird mutation at first on the top part, but then it could have just been like a smaller chromosome or like chromatid or something and they just combined the small.” Donna said: “I’m more confused what the small one and what the tall one is? Like why—like, is this one chromosome I’m looking at? I don’t know. I guess I’m confused of is this how many chromosomes am I looking at? Is this one? Is this two? Is this the same chromosome?”

Participants also produced unexpected interpretations of the labeled alleles. Jimmy interpreted the two lines for each homologous pair as the “two strands, like the double stranded DNA for each chromatid,” and related the capital and lowercase letters from the heterozygous pair of chromosomes as “the uppercase is for the leading strand and the lowercase is for the lagging strand.” The allele “q” was confusing for participants like Donna, who said: “I know there’s a q arm. So I’m going to assume that’s what the two Qs are. And then I thought it was q and p, but I guess not.”

While we focus our analysis here on areas of student misunderstanding to highlight concepts that may need additional instructional scaffolding, we briefly want to emphasize that many participants had productive and insightful ideas about the effectiveness and limitations of the abstract model of chromosomes. Claire (sophomore biotechnology and molecular bioscience major) noted that abstract models are often made when you need to quickly express an idea, such as when making sketches during class: “Sometimes people, will you know, like if they’re sketching on the whiteboard, they’re not going to draw out the full thing, right? They might just draw lines. And drawing the dots helps you kind of see, oh, like ‘top half, bottom half,’ especially if you’re talking about the location of where a certain gene might be on the chromosome—it kind of helps you see it better.” Piper (senior biology secondary education major) emphasized that in addition to the allele labels, the abstract sketch could benefit from additional symbols or notations to emphasize the difference between homologous chromosomes: “I think, like, if I was representing it differently, I’d probably do like maybe a different squiggly line or like a different color—something just because they might get mixed up, especially when people are redrawing them.” By evaluating the abstract model, these participants demonstrated their understanding of the conventions used to represent chromosomes and genes as well as their metacognitive abilities for reflecting on how they or their peers might be thinking about visual representations of chromosomes.

### How do undergraduate students use visual literacy skills to reason about the division of chromosomes during meiosis?

Three-quarters of participants (26 of 35) selected the correct answer to the GCA multiple choice question, indicating an item difficulty on par with what was previously reported (Smith et al., 2008); however, we found that most participants selected the correct answer for the wrong reasons (Table 3). Only 23% of participants (8 of 35) answered the concept assessment question by correctly representing the division of chromosomes during meiosis in their sketches and associated verbal descriptions.

We were surprised to see that 37% of participants (13 of 35) justified their answer using a Punnett square and that 11 of the 13 participants who drew a Punnett square selected the correct answer to the GCA item. Although Punnett squares were the most common way for participants to approach this concept assessment question, there was a wide range in how participants misunderstood the symbols and misapplied the logic of Punnett squares (Figure 7). Participants like Peggy (junior biomedical science major), Leland (junior biology major), and Piper (senior biology secondary education major) recognized that a two-by-two matrix could generate the four different allele combinations present in the correct answer but did not realize that Punnett squares are not appropriate for identifying combinations of alleles from a single parent. Piper initially had inklings that Punnett squares were associated with a single allele, reflected in her drawings of two separate matrixes for the F and Q genes, but rationalized her final answer with a take on a Punnett square that combined both the F and Q genes. Participants who misapplied Punnett squares tended to demonstrate other novice-like ideas, such as Peggy’s conflation of genotype and phenotype, Leland’s inconsistent ordering of alleles inside the matrix, and Piper’s use of the term “monohybrid” to describe a Punnett square for a meiotic event rather than a cross.

**Figure 7.**
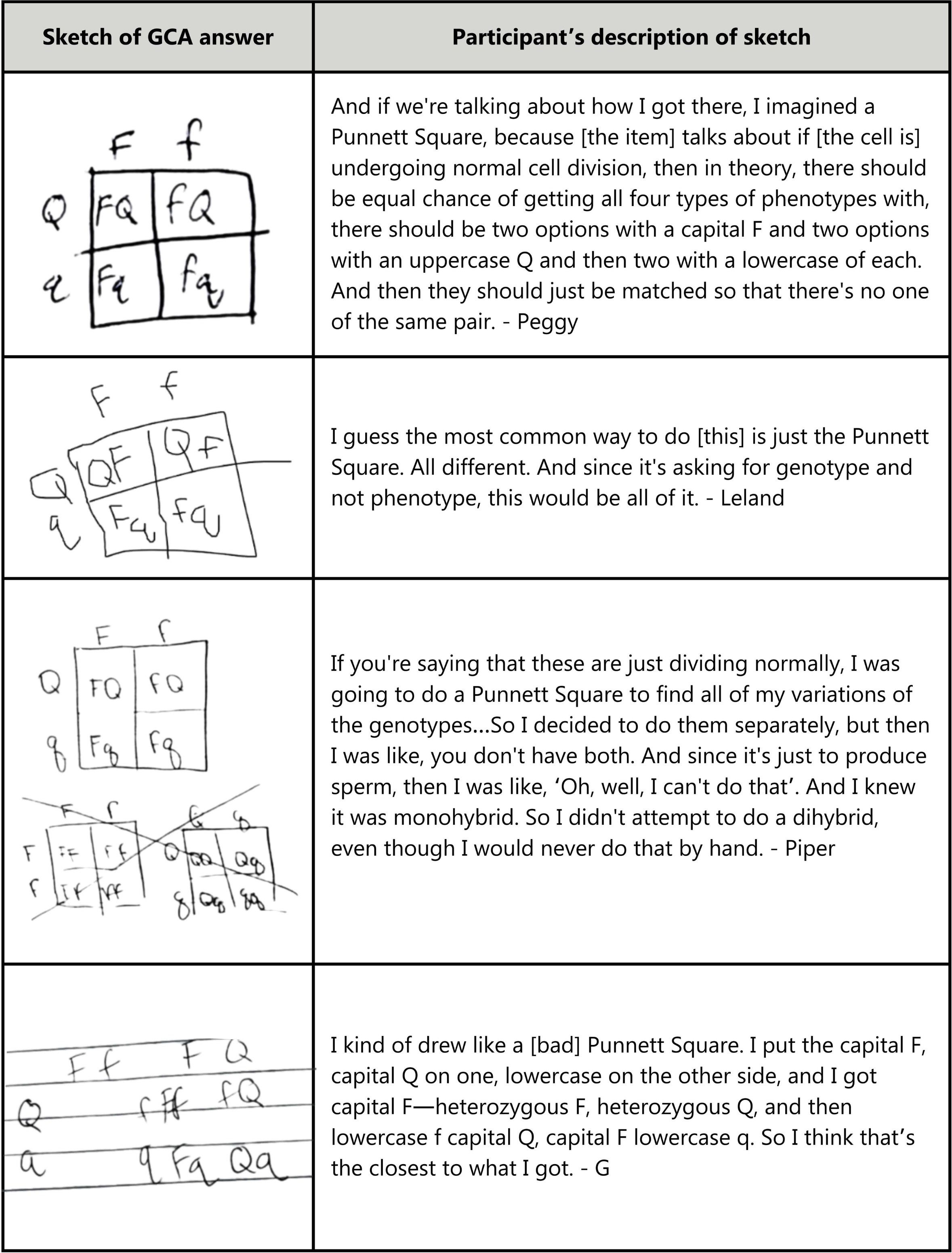
Participant sketches of Punnett squares and associated reasoning to explain their answer to the GCA item. Participants Peggy, Leland, and Piper selected the correct answer to the GCA item. Participant G selected the incorrect answer to the GCA item.

We also saw participants who drew Punnett squares arrive at the incorrect answer to the concept assessment (Figure 7). The participant G (sophomore biotechnology major) drew the structure for two Punnett squares, one which shows F and Q alleles on separate axes (pictured on the left) and one which has a combination of F and Q alleles on each axis (pictured on the right). G then selected the answer which indicated the possible sperm genotypes included Ff, Qq, FQ, fq, Fq, and fQ, and provided the rationale that “I think that’s the closest to what I got” based on their completed Punnett square. The rationale of the “closest to what I got” is interesting here, as the Punnett square only provides four combinations of alleles while the answer G selected contains six possible combinations.

In addition to puzzling uses of Punnett squares, we saw participants arriving at the correct answer for other unexpected reasons (Figure 8). Many times, participants answered thequestion based on their initial (often incorrect) conceptions of chromosomes even though the text from the item states “two pairs of chromosomes.” Heather (sophomore biotechnology and molecular bioscience major) drew the dividing cell with X-shapes consisting of both homologous chromosomes. Each X-shaped structure in Heather’s drawing contains both the dominant and recessive alleles, and she completed meiosis without replicating the chromosomes and with only one cell division. This may be related to Heather’s initial interpretation of the GCA image where she stated “in terms of chromosomes that have already been replicated, there’s two.” Ristan (senior biochemistry and molecular biology major) produced a similar drawing of the crossed homologous chromosomes. During the initial interpretation of the concept assessment image, Ristan said “I’m assuming that later, we’re going to get to the point where the chromosomes start connecting, and we get into sort of these X shapes. So this figure kind of lends itself to that as well, because you can imagine those two sort of coming together, forming that X.” Ristan’s drawing here reflects his initial idea of the “connecting” chromosomes.

**Figure 8.**
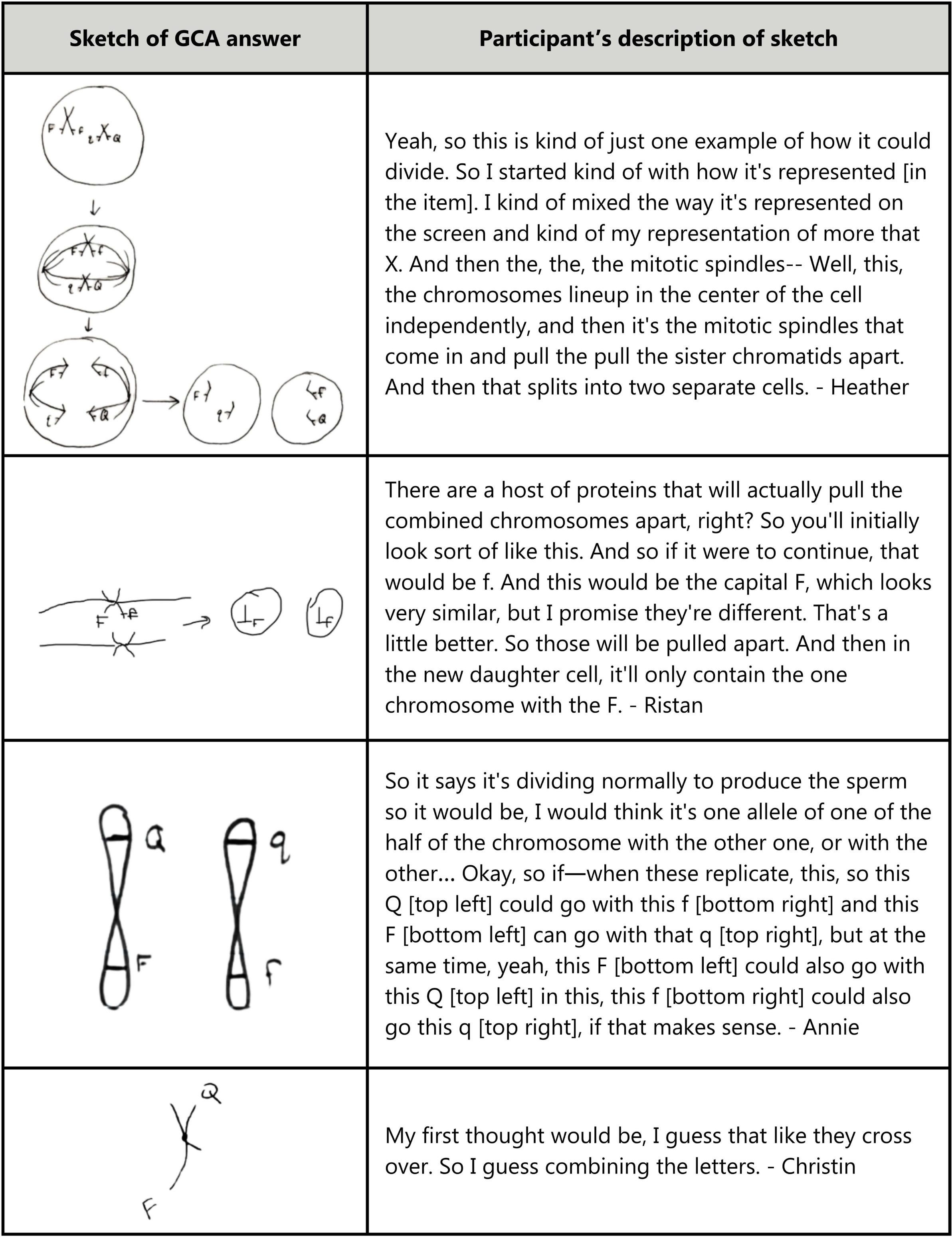
Participant sketches of their answer to the GCA item and associated reasoning to explain their answer. Participants Heather, Ristan, Annie, and Christin selected the correct answer to the GCA item.

We saw more unique interpretations of connecting chromosomes in Annie and Christin’s drawings (Figure 8). Annie only included two chromosomes in her sketch, and the Q and F genes (initially on separate chromosomes) were now depicted on the same chromosome. Annie also used the telling phrase “when these replicate,” indicating that she believed crossing over of alleles occurs during the replication process. We see a similar misunderstanding of crossing over in Christin’s sketch in which she stated that two non-homologous chromosomes are connected at the centromere to represent the process by which chromosomes “cross over.” Notably, both students chose the correct answer choice.

When we asked participants to create sketches to represent their selected answer to the GCA item, we found that many (n = 12) adopted the visual conventions of the GCA item. Even though none of the participants initially represented chromosomes with the I-shaped lines (Figure 3), we saw many students translating their understanding of chromosomes into this representational format in their follow-up sketches. Four of the participants used this shape and style of chromosomes to correctly answer the GCA item for the correct reasons. The remaining eight participants sometimes arrived at the correct answer but their sketches indicated an incomplete or incorrect reasoning, such as failing to replicate the I-shaped chromosomes, resulting in the products of meiosis as only two cells (rather than four) or as containing only a single chromosome (as a result of dividing non-homologous chromosomes in meiosis I and dividing homologous chromosomes during meiosis II).

## DISCUSSION

Chromosome structure and function is foundational knowledge in biology and is necessary to understand the *Vision and Change* core concept of “Information Flow, Exchange, and Storage” (AAAS, 2011). As chromosomes are microscopic and their structural elements (genes, nucleotides) are submicroscopic, biologists often emphasize their relevant features using abstract models. The ability to interpret and use such abstract models are key to *Vision and Change* competencies related to modeling and scientific communication (AAAS, 2011). Despite the importance of these topics in undergraduate education, our findings suggest that chromosomes are parts of introductory biology curricula where college students may need additional instructional support. By probing visual literacy skills, we found a mismatch between declarative knowledge about chromosomes and mental models of chromosomes. Similar to previous research (Newman et al., 2012), our findings here suggest that students’ mental models of chromosomes often need additional refinement to align with how experts typically use abstract models of chromosomes. Our work also mirrors findings in chemistry in which undergraduate students presented with multiple representations of the same concept had difficulties interpreting the main features of abstract models and offered conflicting interpretations across representations (Allred & Bretz, 2019). In this discussion, we highlight particular concepts participants misunderstood and how biology instructors might choose to address these concepts in their teaching.

### Confronting conceptual misunderstandings of chromosomes

While many students were familiar with the mode of representing chromosomes (M in C-R-M), they differed from experts in their conceptual understanding (C) and the ways in which they used their conceptual understanding to reason about the abstract representations (C-R). Here, we focus on where students’ chromosome visual literacy skills were primarily hampered by errors in their conceptual knowledge. For a full list of the conceptual misunderstandings we encountered in this study, see the Supplemental Material.

The most common conceptual misunderstanding was the idea that sister chromatids can be separated and exist as independent structures within the same cell prior to cell division— conflating a sister chromatid with an unreplicated homologous chromosome. This conflation was evident in participants who made the statement that sister chromatids are supposed to be “paired together,” which at face value is a correct statement, but is incorrect when applied as a description of the pair of unreplicated homologous chromosomes in the GCA image. We have previously documented this confusion of homologous chromosomes for sister chromatids (Wright & Newman, 2011), and our finding here bolsters existing evidence that the distinction between chromatids and chromosomes remains a persistent point of conceptual difficulty for students (Clark & Mathis, 2000; Dikmenli, 2010; Kindfield, 1994c).

Similar to how we observed participants identifying homologous chromosomes as “separate” sister chromatids, we also saw the tendency for participants to want to combine two homologous chromosomes into single structure. Previous research has shown that students produced similar representations of “two-DNA-molecule chromosomes” to represent the process of fertilization (Kindfield, 1991). In the previous study, students joined two haploid gametes, each containing a chromosome with the allele “A” or “a,” into a single X-shaped chromosome with both “A” and “a” alleles upon fertilization, but here we saw participants using this “two-DNA-molecule chromosome” representation *at the start of meiosis*.

One way to address some of this confusion about chromatid and chromosome structure is to incorporate active learning opportunities in which students use physical, hand-held models as reasoning tools for visually distinguishing between sister chromatids and homologous chromosomes (Newman & Wright, 2017; Smith & Kindfield, 1999; Wright et al., 2021; Wright & Newman, 2011). Instructors may also consider asking students to reason about the purpose of centromeres in abstract models as a way to clarify the difference between chromatids and chromosomes. Instructors can present multiple models of replicated and unreplicated chromosomes and ask students to determine the total number of chromosomes, emphasizing that biologists often include centromeres in abstract models because counting the number of centromeres is a way to quickly determine the number of chromosomes in a representation.

We were surprised by students who merged non-homologous chromosomes into a single chromosome structure. For example, participants took the two non-homologous chromosomes (denoted with alleles F/f and Q/q), and merged them into a single chromosome containing both F and Q alleles (Figure 8). Interestingly, we provided participants with a representation of chromosomes in a single straight line, but participants that combined homologous chromosomes adopted representations with bowling-pin shapes or curved lines that were not present in the GCA image. It is possible that the students who did not adopt the conventions of the I-shaped line model of chromosomes from the GCA image did not fully understand what the straight lines represented, highlighting the importance of explicitly asking students to evaluate the conventions of chromosome representations when using them as a visual aid when teaching.

Curiously, many participants solved the GCA question by creating Punnett squares to describe the outcomes of one cell undergoing meiosis. These participants were relying on their understanding that Punnett squares are associated with alleles, but Punnett squares are used to predict allele combinations of offspring from a cross between two individuals, not one individual cell undergoing specialized division. While a two-by-two matrix can be a productive strategy for visually sorting combinations when one item in each of two pairs will be part of a combination, referring to the model as a “Punnett square” is problematic. Our finding provides a chain of evidence that students are misapplying Punnett squares in many different genetics contexts (Newman et al., 2021; Shaw et al., 2008; Stewart, 1983; Strand & Boes, 2019). To reinforce the intended purpose of Punnett squares, we suggest that instructors encourage students to make a drawing that indicates each allele on the “outside” of the Punnett square is representative of a sperm or an egg.

We anticipate that many of these conceptual errors result from the phenomenon that chromosome-related concepts are often presented in “twos.” The “twos” occur in the two chromatids within a replicated chromosome, two chromosomes in a pair of homologous chromosomes, and two different pairs of chromosomes (i.e., 2n = 4) that frequently appear in common diagrams of chromosomes (including the one in the GCA). This series of “twos” may in-part explain why participants often conflated two homologous chromosomes in the GCA image for two sister chromatids. To remove one “two” from this challenge, we encourage instructors to use examples with at least 3 pairs of chromosomes (i.e., 2n = 6). Using examples with three pairs of chromosomes may create dissonance with the intuitive ideas that students have about using monohybrid Punnett squares, and may lead to more productive methods of reasoning about the behavior of chromosomes during meiotic events.

### Reckoning with unexpected reasoning about chromosomes

We were surprised at apparent mismatches between participants’ conceptual knowledge about the common modes of representing chromosomes (C-M) and how they reasoned about the meaning of the shapes and symbols in those abstract representations (R-M). We describe several examples here of unexpected ways that participants used shapes and symbols, and provide a full list of the unexpected reasoning we encountered in this study in the Supplemental Material.

One of the most common instances of unexpected reasoning was how several of the participants used loop symbols to indicate that chromosomes were like vessels that could be “filled” with DNA or that each loop was a separate “chunk” of DNA. We speculate that participants’ reasoning about the meaning of the loops may be related their prior experiences and their intuitive sense for symbolism. Multiple participants (Jimmy, Ristan, Ronette, Norma, Andi) described chromosomes as structures that “contain information” or “contain alleles,” and it is possible the colloquial use of the word “contain” may evoke intuitive ideas of chromosomes as “containers” that can be filled. We also saw that some participants thought of separate loops as representing distinct molecules of DNA. Participants described replicated chromosomes as consisting of “four parts” or represented unreplicated chromosomes as two separate “clusters” of DNA separated by the centromere. The misconception that chromosomes contain multiple molecules of DNA has been previously documented (Kindfield, 1991) and was even the subject of an item on the Introductory Cell and Molecular Biology Concept Assessment (Shi et al., 2010), but exactly how we saw participants conceptualizing loops as “chunks” of DNA is a new finding. Our work here provides further support that students may be confused by chromosome structure in ways that hinder their understanding of the nature of the relationship between DNA and chromosomes (Saka et al., 2006). To clarify how loops are used to represent chromosome structure, we recommend that instructors verbally and pictorially explain that each unreplicated chromosome is a single continuous strand of DNA. Instructors may also ask students to work in small groups to compare and contrast models where chromosomes are represented as loops and as lines, particularly using this comparison to probe how students are thinking about what the loops exactly mean.

We highlight another example of unexpected reasoning in the side-by-side chromosomes that several participants drew. Some participants used side-by-side structures to represent a replicated chromosome, while to others these structures symbolized a pair of unreplicated homologous chromosomes. Because the drawings lacked centromeres, it was impossible to know whether the participant was using their drawing to represent one chromosome or two without a verbal explanation. This finding echoes previous work which emphasized the importance of including centromeres in chromosome models as a way to clearly signal the intended number of chromosomes (Smith & Kindfield, 1999). While the purpose of abstraction is to make simplifications, we suggest that abstract chromosome representations should not be simplified to the point of erasing centromeres as a visible feature. As biology students are likely to encounter a variety of chromosome models, we recommend that instructors ask students to evaluate the benefits and limitations of representing chromosomes with and without centromeres. In these instructional exercises, we encourage instructors to be intentional about describing the centromere as a specific segment of DNA within the chromosome to avoid reinforcing the misconception that the centromere is a separate structure or introducing confusion between centromeres and kinetochores.

### Incorporating visual literacy into instruction and assessment

During our interviews, many participants justified their drawings or interpretations of chromosomes by referring to what they had seen in their biology classes or in textbooks. Given the number of conceptual misunderstandings that we encountered, we suggest that there is room to improve how students engage with representations of chromosomes in their classes and course materials. Traditionally, students are likely to have seen prepared graphics in their textbooks and in PowerPoint slides, and this way of presenting graphics and images reinforces an intrinsically passive method of learning via images (Higley et al., 2024; Rotbain et al., 2005; Wright et al., 2022). Teaching using visual representations should go beyond putting an image up on a slide or labeling visual structures. While labeling visual representations may help students memorize structures such as components of a nucleotide in a chemical structure (Rotbain et al., 2005), using visual representations to memorize facts about science is not an end goal. Rather, the *Vision and Change* core competency of “Ability to Communicate and Collaborate with Other Disciplines,” suggests an end goal of science education is for students to be able to *engage with* visual representations and build conceptual models that help them participate in the *processes of science* (American Association for the Advancement of Science, 2011; Clemmons et al., 2020b).

We encourage biology instructors to adopt a more active approach to using visual representations in their courses. We have provided a few sample active learning tasks in this discussion (e.g., asking students to work in small groups to evaluate models). We recommend instructors consult the Visualization Blooming Tool (Arneson & Offerdahl, 2018) for example active learning tasks that can help foster the development of visual literacy skills.

Incorporating visual literacy into instruction does not need to take much class time. Asking students to consider strengths and limitations of a chromosome model requires less than five minutes. In addition to using active learning to teach visual literacy, we encourage instructors to take a few seconds to verbally explain the diagrams, figures, and visuals they use in class. When drawing an X-shaped chromosome, instructors can quickly tell students that the “X” represents a replicated chromosome and circle the two sister chromatids represented within that shape. When drawing a centromere using a circle, instructors can mention that even though they made a separate drawing stroke to create the centromere, it is still part of the same single DNA molecule in that condensed chromosome structure.

While interpreting pre-fabricated visual representations is an important part of biology education, we also encourage both instructors and students to be engaged in creating visual representations during class time. Historically, students are rarely asked to sketch in their science classes (Ainsworth et al., 2011; Van Meter & Garner, 2005), but there is tremendous educational value in drawing (Quillin & Thomas, 2015). Drawing is an important process for biological literacy that is associated with understanding representations, developing scientific reasoning, and effectively communicating science (Ainsworth et al., 2011). While some instructors and students may believe drawing requires a strong artistic talent, we emphasize that sketches do not need to be of the highest artistic quality to be meaningful representations in biology (Quillin & Thomas, 2015). In fact, quickly-constructed “bad” drawings that require interpretation can increase student interaction with the representation and help to reinforce that sketches are useful and acceptable in biology (Higley et al., 2024). These quick sketches that use abstract representations can be powerful learning tools because they distill complexity to show only what is most essential to understand a concept.

In line with principles of Backward Design (Wiggins & McTighe, 2005), instructors who teach visual literacy should also assess visual literacy. Instructors can assess visual literacy with formative assessments, such as evaluating the drawings and explanations that students make on whiteboards during class time, but instructors should also assess visual literacy on summative assessments (e.g., exams) as well. Because content on summative assessments signals to students the knowledge and skills that instructors value and prioritize (Jensen et al., 2014; Scouller, 1998; Uminski et al., 2024), if visual literacy skills are something an instructor wants students to develop, visual literacy skills need to be incorporated into summative assessments. When asking students to draw on an exam is not feasible, assessments may effectively measure student understanding of abstract models if they integrate pre-fabricated models into multiple choice options rather than relying on options that are only based in text. We suggest instructors consult the Drawing-to-Learn framework (Quillin & Thomas, 2015) for a robust set of recommendations about facilitating assessment of visual literacy skills.

Our work underscores the importance of valid and reliable visual literacy assessments. We found that the majority of participants in our study selected the correct answer to the GCA item for the wrong reasons. By asking them to create a visual representation of their answer to the GCA item, we uncovered that the majority of participants had incomplete or incorrect mental models of chromosomes or meiosis and were getting the item right even though they had an incomplete understanding of the content the item was designed to assess. Had we relied on the multiple-choice answer alone, we would have vastly overestimated our participants’ true conceptual understanding of chromosomes and meiosis. Our finding here emphasizes both the limitations of traditional multiple-choice assessments in which students can select the right answer for the wrong reasons (Cerchiara et al., 2019; Fisher et al., 2011; Fulmer et al., 2015; Haudek et al., 2012) and the importance of assessing students’ visual literacy skills as a robust method of gauging conceptual understanding (Arneson & Offerdahl, 2023; Offerdahl et al., 2017; Schönborn & Anderson, 2006). We emphasize the need for biology education researchers to develop valid and reliable assessment instruments that specifically measure visual literacy skills in biology.

### The DNA Landscape provides a framework for visual literacy in molecular biology

We highlight the DNA Landscape (Wright et al., 2022) as a useful framework for instructors looking to teach visual literacy skills and for researchers investigating visual literacy in the context of molecular biology. For example, we found evidence of student misunderstanding of chromosomes when their drawings mixed abstract representational conventions. We observed that participants who drew representations of chromosomes that mixed the abstract X-shapes with the more realistic string representations (Figures 4 and 5) tended to have incorrect ideas about chromosome structure and function, often confusing how more experienced biologists traditionally use the string representation. Identifying where students are mixing representations within columns of the DNA Landscape can provide insights into where students may be facing conceptual challenges in molecular biology.

We suggest that instructors use the DNA Landscape to inform their choices related to *what* and *how* they are visually representing molecular biology concepts. For example, instructors teaching about chromosomes should attend to asking students to interpret multiple representations within the same column, making sure to clarify that chromosomes can be represented using abstract lines, as maps or ideograms, or more realistically as strings. In addition to providing opportunities for students to reason with multiple representations within a column of the DNA Landscape, we encourage instructors to incorporate small group discussion and ask students discuss with each other why biologists might use one representation over another. We also suggest instructors take a moment of class time to ask students to reason across the columns of the DNA Landscape to help illuminate the relationships between nucleotides, genes, and chromosomes. Instructors can ask students to consider why it is inappropriate to use a Punnett square to answer a question about meiosis or to think about the relationship between the double helix and chromosome structures. Instructors might ask students if the helix “fills” the chromosome, as some of our participants here have suggested. By giving students opportunities to grapple with multiple representations across scale and abstraction in the DNA Landscape, instructors can help their students develop the fluency in visual literacy that is necessary for deep understanding of the foundational concepts in molecular biology.

Biology education researchers can use the DNA Landscape as a framework for designing research studies and assessment tasks. Researchers may design future work to study how biology students are thinking about the similarities and differences between representations within columns of the DNA Landscape. There are also opportunities for future work to study how students are conceptualizing scalar relationships across the columns of the DNA Landscape. When developing research studies or assessment tasks related to visual literacy in molecular biology, we suggest researchers consider incorporating representations across the DNA Landscape.

### Future directions

We found that misunderstandings of chromosome structure and function are “sticky,” often persisting in students who have already entered advanced undergraduate coursework in their third and fourth year of college. Our findings emphasize areas for future research to investigate where students may have derived some of their incorrect ideas and misconceptions about chromosomes. We anticipate that some of these misunderstandings we observed may be related to the colloquial ways that students see chromosomes discussed and represented in K-12 life science curricular materials (Cho et al., 1985). We highlight the use of the word “half” to describe chromatids as one example of this imprecise language that we suspect created confusion in students’ understanding of chromosomes. Multiple students referred to sister chromatids (or, in a few cases, a pair of unreplicated homologous chromosomes) as being two “halves” of a replicated chromosome. While our participants frequently used the word “half,” and the term “half” appears in open education resources about genetics geared towards a K-12 audience (National Human Genome Research Institute, 2024), this term was never used to describe chromatids in undergraduate biology textbooks (Brooker, 2011; Freeman et al., 2017; Lodish, 2008; Morris et al., 2022; Solomon et al., 2015; Urry et al., 2016). Learning theory suggests that schemas are established based on initial learning and these schemas can be difficult to change (Piaget, 1952); thus the incomplete ideas about chromosomes that are assimilated during middle and high school can persist even when more correct information is presented in undergraduate courses. As the first learning about chromosomes can shape what is understood (or misunderstood) in future learning, we recommend further investigation into the representations of chromosomes in K-12 contexts.

We also recommend future research efforts to uncover additional insights into the types of representations, learning activities, and assessments that are most beneficial to the development of visual literacy skills across a variety of molecular biology contexts. Our work leaves questions about how different types of models in molecular biology may help or hinder student understanding of particular concepts. While it was not a focus of our research, we found that that many students mentioned seeing representations of chromosomes in their classes and textbooks, but no students mentioned seeing such models on exams or on other class assignments. This leads to questions as to how students are engaging with the abstract representations in these in-class, out-of-class, and assessment contexts. There are remaining questions about how is learning in molecular biology is affected by the ways students engage with models during class time, impacted by the frequency with which models are used in a course, and influenced by how models are integrated into summative assessments.

### Limitations

We necessarily focus our results on the errors in student conceptualizations of chromosomes, as these errors provide novel insights into student thinking that may be useful for informing instruction and further education research on conceptual understanding in molecular biology.

We recruited participants from two institutions. It is possible that biology instructors at other institutions may emphasize different aspects when teaching about chromosomes, and as such, we do not make claims that the misunderstandings we observed in our sample population are representative of what misunderstandings might be present in other student populations.

We informed the participants in our study prior to the interview that they would be asked to draw during the interview protocol. As a student’s attitude, self-efficacy, and interest in drawing biological models can affect their use and engagement with models (Quillin & Thomas, 2015), we likely only recruited students who already had a positive attitude, high self-efficacy, and high interest in drawing biological models. Given this self-selection bias in our recruitment and sampling, we highlight the need of additional visual literacy research conducted in ways that can capture a wider range of student affect towards drawing.

## CONCLUSION

Understanding chromosomes is necessary for understanding more complex concepts in genetics. Even though the notion of chromosomes begins as early as middle school, it remains a difficult concept for many undergraduate biology students, which may be tied to their incomplete or incorrect mental models of chromosome structure and function. These difficulties may not be evident in students’ verbal descriptions of chromosomes and are only revealed when probing students’ visual literacy skills. Asking students to create drawings and interpret visual representations provides key insights into students’ conceptualizations of chromosomes. We encourage biology instructors to incorporate opportunities to develop students’ visual literacy skills in molecular biology by using multiple representations of chromosomes at different levels of abstraction, engaging students in sketching representations during class, and asking students to evaluate conventions used in visual representations.

## Acknowledgements

Funding provided by NSF DGE 2222337. We thank Christian Cammarota, Meredith Michetti, Shreya Sujith, Aneesh Nallani and Jupiter Chen for help validating the codebook. We are grateful to the research participants for sharing their ideas with us. We thank Michelle Smith and Jenny Knight for their permission to publish a reproduction of a figure from The Genetics Concept Assessment.

