## Supplemental Material for "Probing visual literacy skills reveals unexpected student conceptions of chromosomes"

| Misunderstanding | Example quotes | Example sketch | Percent of participants |
|---|---|---|---|
| Sister chromatids can be separated and exist as independent structures within the same cell (i.e., conflating sister chromatids for homologous chromosomes) | "The chromatids aren't paired together?" (G)<br><br>"The sister chromatids or whatever aren't connected at all – Like it looks like they're two standalone" (Alex)<br><br>"There's two strands and there's two centrosomes, and typically, I think they're kind of fused together" (J)<br><br>"I usually imagine them like connected like I did…because they're like separate it might be something that I thought was a little confusing at first" (Quentin)<br><br>"I would assume it's maybe two halves of the chromosome just pulled apart. Instead of being in that X, they're just portrayed separately" (Eileen)<br><br>"I basically understood that like, okay, they come together, and then they drift apart, like they got pulled apart" (Sylvia)<br><br>"I think it does [well] like showing that there's two separate chromatids" (Jimmy) | 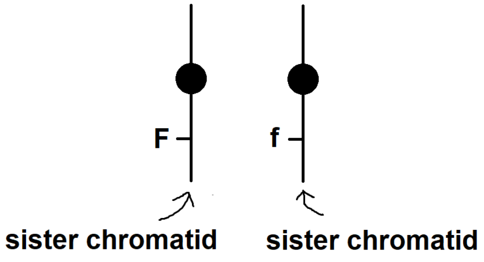<br>(Figure created to illustrate student thinking) | 43% |

| | |
|---|---|
| | "Those are chromatids there, aren't they? So I think that's what's confusing me – the two chromatids come together and make a chromosome" (Claire)<br><br>"I liked that the centromere kind of stays with the side of each sister chromatid" (Maddy)<br><br>"It better shows how the chromatids are split up during mitosis" (Socks)<br><br>"I like how it clearly identifies like the two halves—the two chromatids" (Nadine)<br><br>"I would think like mine shows that they're kind of paired together that they are actually bound together, whereas here, it's kind of showing them as being separated from each other" (Leo)<br><br>"I'm assuming that later, we're going to get to the point where the chromosomes start connecting and we get sort of these X shapes, so this figure kind of lends itself to that as well because you can imagine those two sort of coming together forming that X" (Ristan)<br><br>"You have the little dots, because that's where they used to be connected…We have the chromosome and it's pulled into one pair" (Pluto) |

| | | | |
|---|---|---|---|
| Punnett squares can be used to model meiosis | "If you're saying that these are just dividing normally, I was going to do a Punnett Square to find all of my variations of the genotypes" (Piper)<br><br>"Okay, if it divides normally to produce sperm, the way I see it is that each of the chromosomes will be divided in half, and then equally separated. So then you have to do one of those like Mendelian Punnett squares" (Josie)<br><br>"I imagined a Punnett Square, because it talks about if it's undergoing normal cell division, then in theory, there should be equal chance of getting all four types of phenotypes. There should be two options with a capital F and two options with an uppercase Q and then two options with a lowercase of each. And then they should just be matched that there's no one of the same pair" (Peggy) | 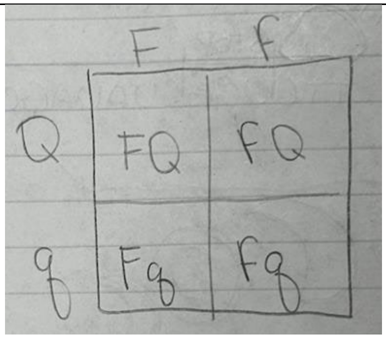<br>(Piper) | 40% |
| There can be different chromatids for the same chromosome | "You can see that each sister chromatid looks pretty much exactly like the other one, and it very clearly demonstrates having different alleles on the sister chromatids" (Socks)<br><br>"I mean, it has all the information needed, it has two different chromatids." (G)<br><br>"One section of the code will be on this end and one section will still be on that side. It's double…for example, if you are homozygous, it'd | 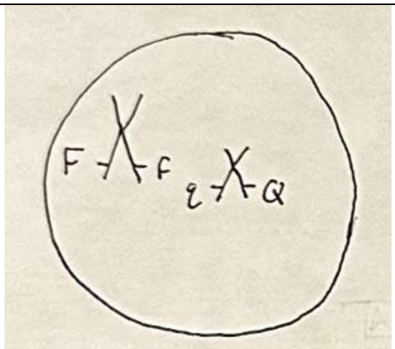<br>(Heather) | 17% |

| | | | |
|---|---|---|---|
| | be like your big E and then like the little e" (Annie)<br><br>"[The letters], they're just like to specify that they're the same chromosome but different sister chromatids" (Maddy)<br><br>"A chromosome – you get one from both your parents – one chromatid from both your parents to make your chromosomes" (Nadine) | | |
| Meiosis occurs with a single cell division | "Because sperm is haploid, to get one half of each of the chromosomes in the original cell" (20, Maddy)<br><br>"So with it being split apart, you'd assume that the two cells will give ou the "captial F, lowercase f" and the "lowercase f, capital F" genotype, and then the two Q variants" (Eileen) | 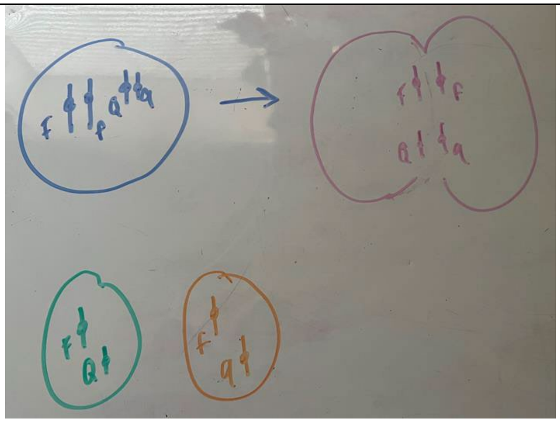<br>(Maddy) | 14% |
| Crossing over can happen between non-homologous chromosomes | "My first thought would be, I guess that they like cross over. So I guess combining the letters" (Christin)<br><br>"So since there's two different alleles for each gene, that would show like these two would be shown there at FF, and there would be a crossover with the Q to form different variants" (Jimmy) | 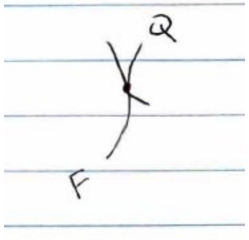 <br>(Christin) | 11% |

| | | | |
|---|---|---|---|
| | "It says it's dividing normally to produce the sperm, so I would think it's one allele of one of the half of the chromosome with the other one" (Annie)<br><br>"It's like two chromosomes, so F and the Qs together, yeah. And then you have the exchange of genetic information" (J) | | |
| Replicated chromosomes consist of four distinct parts | "I always thought it was made up into like four little blocks. Like the top block was a completely different color from the bottom block and then it flipped on the other side" (13, Eileen)<br><br>"I believe [each of the 4 loops] is all like piece of the genetic code. I mostly know what they do in like mitosis, when it's like that determination and like cell splitting, stuff like that, because they split off and they all contain a little bit of information that is used to help make the genetic code or to be copied" (Ronette)<br><br>"I'm assuming that the entirety of the chromosome is encapsulated in the circle, and that the capital and lowercase letters could represent the different of the—like the four quadrants that I drew" (Peggy) | 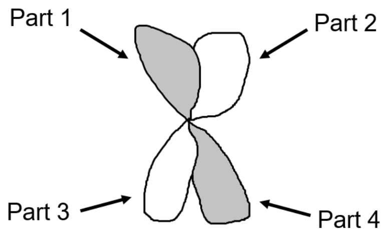<br>(Figure created to illustrate student thinking) | 11% |

| | | | |
|---|---|---|---|
| Non-homologous chromosomes separate during meiosis II | "Each of the four daughter cells will only receive informaiton from one sister chromatid per chromosome pair, so only one allele of each type" (Socks)<br><br>"I know that we produce like four, like you would get four individual cells from it with one, and they would be haploid. So it couldn't be, it couldn't have two alleles in it" (Leo) | 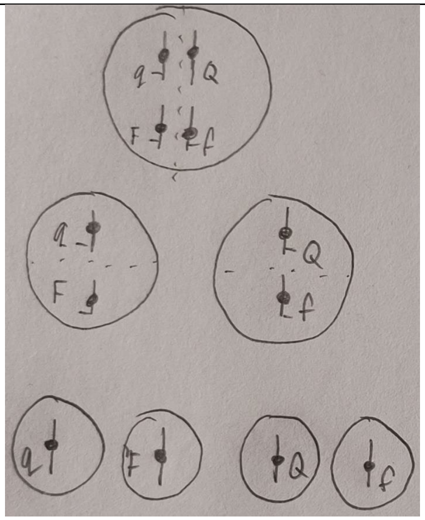<br>(Socks) | 11% |
| The dot that represents the centromere is a gene | "I'm guessing the line is a chromosome, the circle's maybe that's just a highlighted region of like where the gene is" (Leo) | 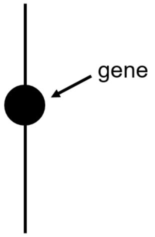<br><br>(Figure created to illustrate student thinking) | 9% |
| Centromeres do not contain genetic information | "Like you have the centromere in the middle…or like the DNA that would be on both sides of it" (Quentin)<br><br>"Circle's the centromere and loops is just the genetic sequence" (Lucy) | 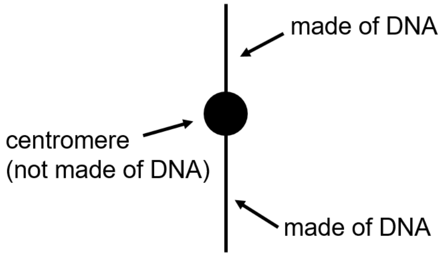<br><br>(Figure created to illustrate student thinking) | 9% |

| | | | |
|---|---|---|---|
| Centromeres can bind homologous chromosomes | "I think the dot is more showing the location of being like this is a connector, but since like the location is like the same on both of them, like it can kind of like connect together –they're supposed to be connected together" (Sylvia)<br><br>"One of them is really tiny and one of them is very big. I'm not quite sure why that's a thing, unless they're different types of chromosomes and they have the little dots on the center, which is maybe where they would like connect?" (Regina) | 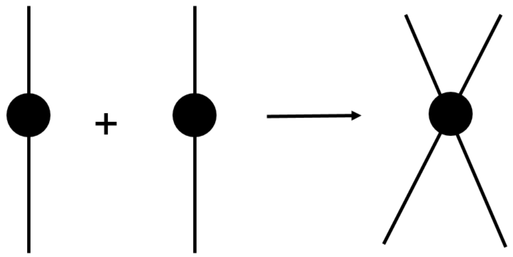<br>(Figure created to illustrate student thinking) | 6% |
| There are separate "clusters" of DNA above and below the centromere | "This is like the clusters of the DNA, big clusters…And then this is also another cluster of like DNA" (Teresa)<br><br>"Like you have the centromere in the middle…or like the DNA that would be on both sides of it" (Quentin) | 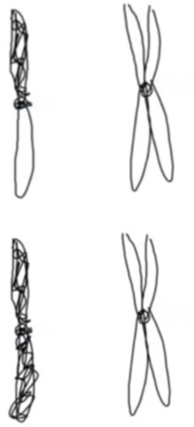<br>(Theresa) | 6% |
| Chromosomes are "part of genes" | "It's like a chromosome was basicially, I thikn it's like part of like genes" (Mia)<br><br>"A chromosome is a part of the DNA that can lead to a gene" (Evelyn) | N/A | 6% |

| | | | |
|---|---|---|---|
| Letters represent labels to distinguish chromosomes (not to distinguish genes or alleles) | "There's probably like two pairs of chromosomes, like each labeled with like, it's own—like one is an F chromosome and the other's like a Q… So it's just like showing you like this is chromosome A and that's like chromosome B" (Mia)<br><br>"I see the Q and I think of the p and q ends of the chromosomes, but I'm not sure about the F. I'm assuming this is a way of labeling the chromosomes though" (Teresa) | 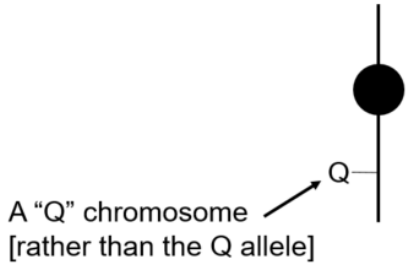<br>(Figure created to illustrate student thinking) | 6% |
| Two parallel lines represent individual DNA strands in DNA replication | "Like the two lines for each one, like there's like two lines on the F and two lines on the Q, maybe that's like the two strands, like the double stranded DNA for each chromatid. I guess like maybe the uppercase is for the leading strand and the lowercase is for the lagging strand" (Jimmy)<br><br>"I've maybe seen like two lines when you're doing like DNA replication or something, and then it's like you start with two, then you have like so much more, like it's replicating" (Mia) | 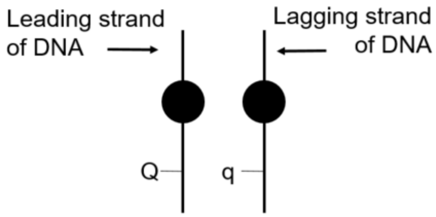<br>(Figure created to illustrate student thinking) | 6% |
| Chromosomes consist of numerous strands of DNA | "I think that's to sort of show that it's not like one strand…that it is many, many strands of DNA all compressed on top of each other" (Ristan) | 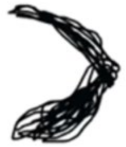<br>(Ristan) | 3% |
| A eukaryotic chromosome contains all of the DNA in an | "They're like the encapsulation of all your DNA" (Jimmy) | N/A | 3% |

| | | | |
|---|---|---|---|
| organisms' genome | | | |
| Chromosomes are only in eukaryotic cells | "I think a chromosome is like, I think it's in eukaryotic cells, and like, or it can also be in plants and animals" (Mia) | N/A | 3% |
| Chromosomes are always "lined up" | "When I think of a chromosome, I think of like 23 lined up when you're doing the one—I can't remember what it's called—but we're looking at the chromosomes like the condensed form and that's that I imagined" (Quentin) | [We presume that this student was referencing a karyotype] | 3% |
| The length of chromosomes is associated with dominance | "Could the longer strands mean it's dominant?" (Annie) | 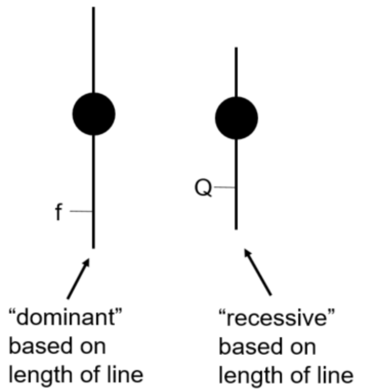<br>(Figure created to illustrate student thinking) | 3% |
| The same genes are located at identical points the same distance above and below the centromere | "There's like a center point of the chromosome, and then from there, it's sort of reflected. So assuming my understanding is correct, you would also expect a similar gene here, that would be like an uppercase F" (Ristan) | 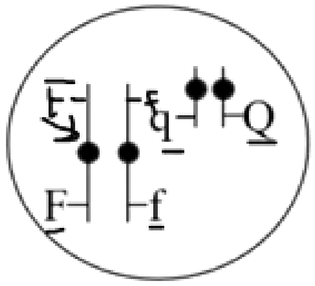<br>(Ristan) | 3% |

| | | | |
|---|---|---|---|
| Longer chromosomes can be composed of smaller chromosomes combined together | "My first thought was that there was like a weird mutation at first on the top part, but then like it could have just been a smaller chromosome or like chromatid or something and they just combined the small" (Brittany) | N/A | 3% |
| DNA can become single stranded through biological processes | "The DNA can be double stranded, and then if it goes through like the other processes, it could become single stranded and whatnot" (Norma) | N/A | 3% |
| Centromeres are separate structures apart from chromosomes or chromatids | "I drew it condensed and I drew it with the sister chromatid with the centromere binding the two pieces together" (Alex) | 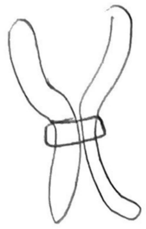 (Alex) | 3% |
